# Bladder mucosal afferents detect UTI and aid pathogen clearance

**DOI:** 10.1101/2025.10.23.684249

**Authors:** Cindy Tay, Harman Sharma, Stewart Ramsay, Georgia Bourlotos, Sarah K Manning, Natalie E Stevens, Feargal J Ryan, Geraint B Rogers, David J Lynn, Andrea M Harrington, Vladimir Zagorodnyuk, Steven L Taylor, Luke Grundy

## Abstract

The bladder is innervated by a complex network of sensory neurons that detect and transmit mechanical and chemical signals to the central nervous system, regulating both urine storage and voiding, as well as mediating sensations of bladder fullness and pain. While stretch-sensitive afferents within the bladder wall are known to monitor filling, the contribution of stretch-insensitive afferents that reside within the bladder mucosa to bladder sensation and function remains unclear. Here, we establish a novel mouse model of selective bladder mucosal afferent denervation using intravesical instillation of resiniferatoxin (RTX) to define the functional roles of these fibres in health and disease. We assessed the impact of mucosal denervation on sensory nerve activity and bladder function in healthy mice and in a model of urinary tract infection (UTI) induced by acute bacterial challenge with uropathogenic *E. coli*. We show that mucosa-innervating afferents do not influence bladder distension responses or voiding function under healthy conditions. In contrast, during UTI, these afferents become hypersensitive and serve as critical drivers of infection-induced pelvic pain and urinary frequency, responses that contribute to enhanced bacterial clearance. Together, these findings identify bladder mucosal afferents as key sensors of pathogenic challenge and reveal a previously unrecognised mechanism that drives behavioural responses to reduce the burden and extent of UTI.

## Introduction

The bladder is innervated by a complex network of sensory nerves with functionally diverse properties that allow the detection of both mechanical and chemical stimuli (1–5). Bladder-innervating afferents have cell bodies within the dorsal root ganglia (DRG) and terminate within the dorsal horn of the spinal cord where they feed into the central nervous system circuits responsible for regulating bladder sensation and function (6–8). Around 75-80% of all bladder afferent endings are found within the detrusor smooth muscle where they are responsible for detecting the degree of bladder wall stretch (4, 5, 9) and transducing sensory signals to the brain to elicit sensations of bladder fullness necessary to support normal bladder function and maintain homeostasis (4, 6, 10, 11). The remaining 20-25% of afferents innervating the bladder have endings closer to the bladder lumen, within the bladder mucosa, and a significant proportion of these exhibit no response to physiological levels of bladder stretch (4, 5, 12–14). The role of these stretch-insensitive mucosal afferents in bladder function has yet to be determined. However, their greater chemosensitivity (4, 13, 15) and location within the superficial layers of the bladder where infection and inflammation predominate (16), has led us to hypothesise that they may serve a crucial role in detecting and signalling the presence of pathological stimuli.

Urinary tract infections (UTIs) are one of the world’s most prevalent infections and the most common pathophysiological challenge to the bladder (17). UTIs are characterised by the development of irritative and painful sensations that are highly distinct from those experienced during healthy bladder function, including pelvic pressure/pain, dysuria (pain/burning sensation when urinating), and severe urgency that drive an increase in urinary frequency (1, 2, 8, 18). In addition to alerting the host to the presence of bladder pathophysiology, these symptoms have also been proposed as important regulators of pathogen clearance by driving an increase in the frequency of urination (8, 19, 20). However, it remains unclear if specific subtypes of bladder afferents are responsible for signalling distinct aspects of bladder sensation to drive these sensory and behavioural responses.

Here we address this knowledge gap by developing a novel mouse model of bladder mucosal afferent denervation utilising the neurotoxin resiniferatoxin (RTX) and assessing the consequences on sensory nerve fibre activity, bladder function and bacterial persistence during acute bacterial challenge. Our results implicate mucosal afferents as key players in the detection of pathological challenge in the bladder and the initiation of urinary frequency to reduce the burden and extent of acute bladder infection.

## Results

### Intra-bladder resiniferatoxin selectively denervates mucosal afferents

Understanding the contribution of mucosal afferents to bladder sensation and function has been hindered by a lack of *in vivo* models to effectively and specifically isolate their function. To address this challenge, we leveraged the potent neurodegenerative properties of resiniferatoxin (RTX) (21–23), the known expression of TRPV1 within bladder afferents (24, 25), and the ability to instil compounds within the bladder lumen following urethral catheterisation to selectively denervate TRPV1-expressing neurons in the bladder mucosa (**Fig 1**)

**Figure 1:**
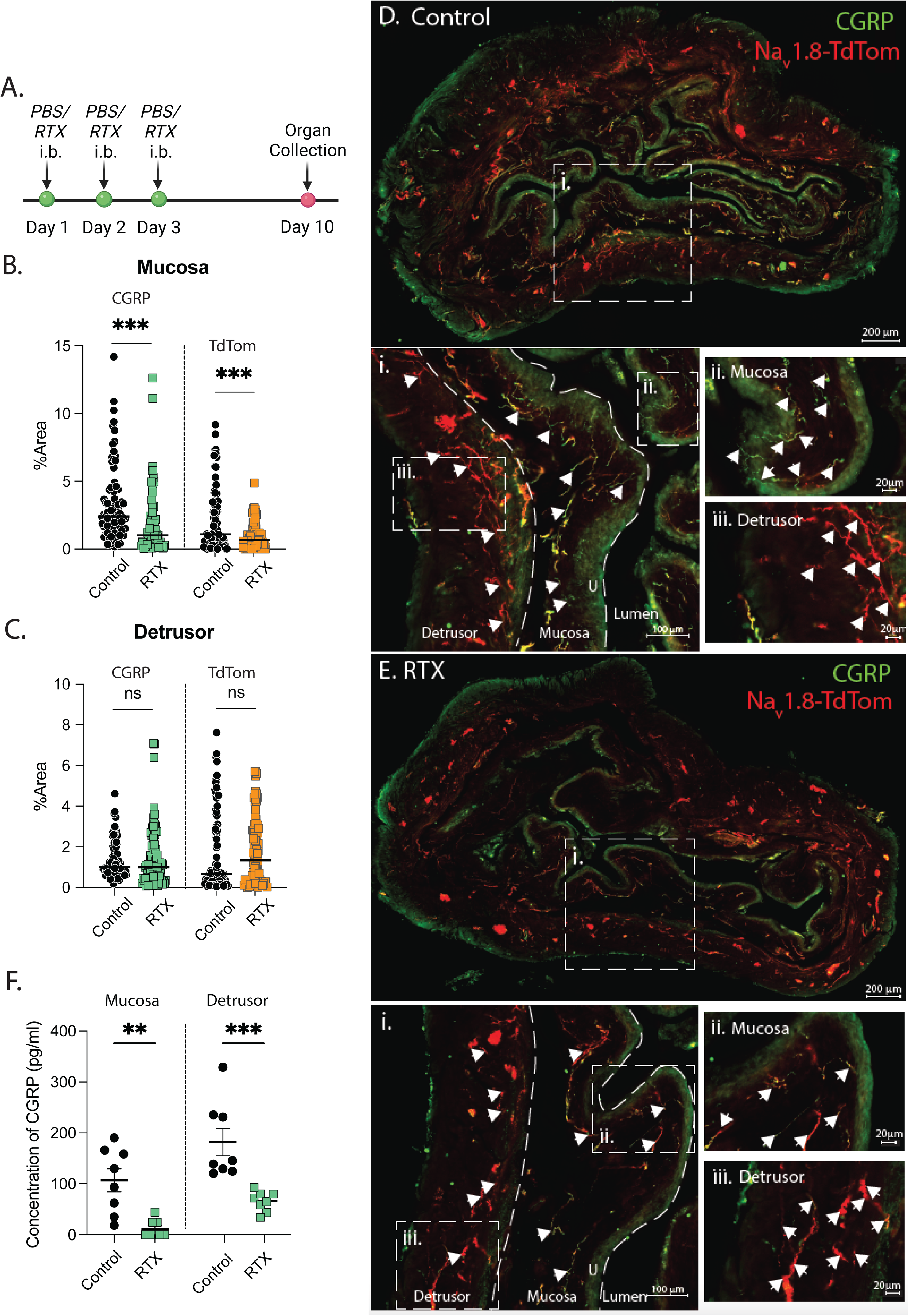
Intravesical RTX reduces the density of CGRP-immunoreactive and Nav1.8-tdtomato expressing fibres in the bladder mucosa. **(A)** Schematic representation of the experimental design for the murine mucosal denervation model. Mice received either RTX or PBS (SHAM) for 3 consecutive days and bladders were assessed 7 days after the final infusion (**B, C**) Quantitative data from cross-sections of bladders showing % Area of the mucosa (**B**) and the detrusor (**C**) covered by Nav1.8-TdTom and CGRP-AF488 fluorescence (n = 90 sections from N = 5 mice/experimental group). Representative photomicrographs showing the distribution of CGRP-immunoreactive fibres (green) and Na_v_1.8-tdTomato expressing fibres (red) in cross-sections (40µm) of the bladder of control **(D)** and RTX-treated **(E)** mice. (**Di, Ei**) High magnification of the bladder wall showing CGRP-immunoreactive and Na_v_1.8-tdTomato expressing fibres innervating the detrusor, lamina propria and urothelium (Mucosa). Inserts from (Di, Ei) show CGRP-immunoreactive and Na_v_1.8-tdTomato expressing fibres within the mucosa (**Dii, Eii**), and detrusor (**Diii, Eiii**) at higher magnification. **(F)** Quantification of the protein concentration of CGRP between control and RTX treated bladders (N=8/experimental group). Data presented as mean ± standard error of the mean (SEM). ^ns^ P > 0.05, **p < 0.01, ***p < 0.001. Statistical differences were determined using non-parametric Mann-Whitney test. Data in F analysed by Welch’s t-test.

Throughout the bladder, sensory fibres labelled by CGRP⁺ and TdTom⁺ fluorescence were identified, forming a dense network innervating the detrusor smooth muscle and mucosa (lamina propria + urothelium) (**Fig. 1B–E, Fig. S1–3**). Intravesical RTX instillation markedly reduced the density of these innervation markers in the bladder mucosa (**Fig. 1B, 1Dii, 1Eii**), but not in the detrusor (**Fig. 1C, 1Diii, 1Eiii**) indicating selective denervation of mucosal afferents. Analysis of distinct regions of the bladder wall (Neck, Middle, Dome) revealed consistent and specific loss of CGRP fluorescence in the mucosa (**Fig S1-2**). Quantification of CGRP concentration within excised mucosa and detrusor tissue showed almost complete abolishment of CGRP in the mucosa, and a significant decrease in CGRP within the detrusor (**Fig 1F**). The effect of RTX on TdTom^+^ fluorescence was greatest within the mucosa of the bladder dome (**Fig S1, S3**). Loss of mucosal afferents occurred in the absence of obvious changes in bladder structure (**Fig S4**), or resident immune cell populations (**Fig S5**), confirming the specificity of RTX in targeting sensory innervation and rapid recovery of any potential neurogenic inflammation induced by RTX treatments (26).

The functional impacts of intravesical instillation of RTX on bladder sensory signalling were assessed through changes in mechanosensory responses of the major bladder afferent subtypes including muscular, muscular-mucosal, and mucosal afferents (4, 5) (**Fig 2**). Aligning with our anatomical data (**Fig 1**), mucosal afferents, which do not respond to bladder stretch but can be activated by light urothelial stroking(4, 5), were mostly absent from RTX-treated electrophysiology preparations (**Fig 2Bi**). Of the mucosal afferents that retained activity, both spontaneous firing frequency (**Fig 2Bii)** and mechanosensory responses to stroking were significantly reduced compared to sham preparations (**Fig 2Biii-iv**). The number of functionally responsive muscular-mucosal afferents, which respond to both mucosal stroking and bladder stretch and have endings that extend throughout the mucosa and detrusor, were also significantly reduced following RTX treatment **(Fig 2Ci)**. RTX-treatment had no significant impact on spontaneous firing frequency (**Fig 2Cii**) or mechanosensory responses to either mucosal stroking **(Fig 2Ciii-iv)** or bladder stretch **(Fig 2Cv-vi)** of remaining muscular-mucosal afferents. In contrast to effects on mucosal and muscular-mucosal afferents, RTX treatment had no significant impact on the number of muscular afferents per preparation **(Fig 2Di)**, spontaneous firing frequency **(Fig 2Dii)**, or mechanosensory responses to both low- and high-intensity load (1-5g) induced stretch **(Fig 2Diii-iv)**. Taken together these data indicate that RTX treatment potently and specifically reduces innervation and function of afferents with endings in the bladder mucosa without major impacts on stretch-sensitive muscular afferents innervating the detrusor smooth muscle.

**Figure 2:**
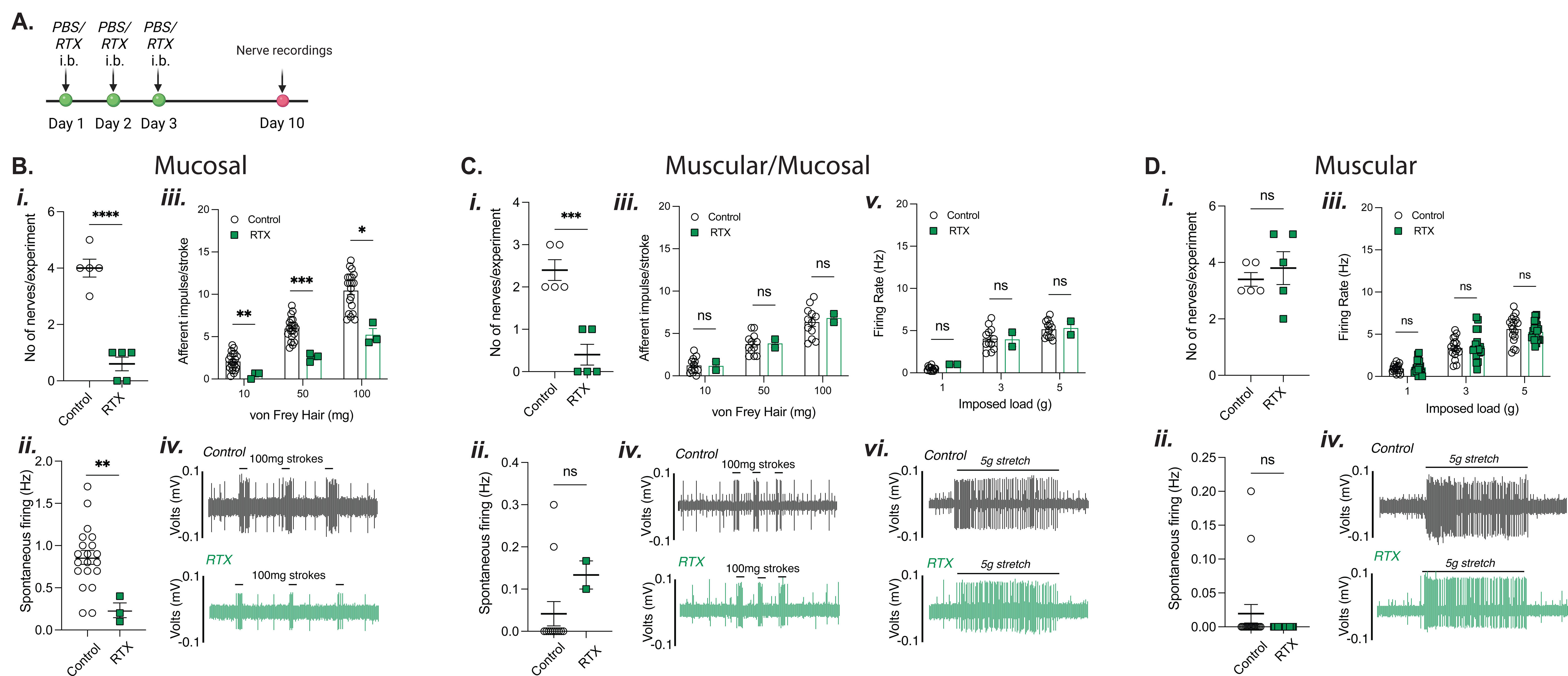
Intravesical RTX reduces mechanosensitivity of mucosa innervating afferents. (**A**) Schematic representation of the experimental design. Mice received either RTX or PBS (SHAM) for 3 consecutive days and bladders were assessed 7 days after the final infusion (**B**) **Mucosal afferents**. (**Bi**) The number of functional mucosal afferents identified from each independent experiment (N=5/group). (**Bii**) Mucosal spontaneous firing frequency (n=20-3 responsive units from N=5 mice/group). **(Biii)** Mucosal afferent responses to von Frey Hair stroking (10, 50, 100 mg) in remaining mucosal afferents. (**Biv**) Representative mucosal afferent nerve profiles of control (black) and RTX (green) treated bladders to 100mg mucosal stroking. (**C**) **Muscular-mucosal afferents**. (**Ci**) The number of muscular/mucosal afferents identified from N=5 independent experiments/group. (**Cii**) Spontaneous firing frequency in remaining muscular-mucosal afferents (n=20-3 responsive units from N=5 mice/group). **(Ciii)** Muscular-Mucosal afferent responses to von Frey Hair stroking (10, 50, 100 mg), and (**Civ**) representative response profiles of control (black) and RTX (green) treated bladders to 100 mg mucosal stroking. (**Cv**) Muscular-Mucosal afferent responses to bladder stretch by imposed load (1, 3, 5 g), and representative response profiles of control (black) and RTX (green) treated bladders to 5g bladder stretch (**Cvi**). **(D) Muscular afferents**. (**Di**) The number of muscular afferents identified from individual experiments (N=5). (**Dii**) Spontaneous firing frequency of muscular afferents (n=17-19 responsive units from N=5 mice/group)**. (Diii)** Muscular afferent responses to bladder stretch by imposed load (1, 3, 5 g) and (**Div**) Representative response profiles of **(i)** control (black) and **(ii)** RTX (green) treated bladders to 5g bladder stretch. Data is presented as mean ± SEM. ^ns^ P > 0.05, * p < 0.05, **p < 0.01, ***p < 0.001, and ****p < 0.0001. Statistical differences for Ai, Aii, Bi, Bii, Ci, Cii were determined by unpaired Mann-Whitney test. For Aiii, Biii, Bv, Ciii, statistical differences were determined by two-way ANOVA with Geisser-Greenhouse correction and Šídák multiple comparisons.

### Denervation of mucosal afferents has no impact on normal bladder sensation and function

Awareness of bladder fullness is provided by integration of mechanosensory signals from the bladder into the central micturition circuits that regulate bladder filling and emptying and is essential for normal physiological functioning of the bladder (6–8). Our RTX-model of mucosal denervation provided a unique opportunity to explore the contribution of mucosal afferents to bladder sensory signalling during bladder filling in an intact bladder (**Fig 3B-D**). Using an *ex-vivo* whole bladder afferent recording preparation we observed robust afferent responses to graded bladder distension (0-50mmHg) in both RTX and sham treated bladders (**Fig 3Bi-ii**), with no significant difference in afferent response to distension (**Fig 3Bi-ii)** and no change in bladder compliance (**Fig 3C**) following RTX treatment. Subsequent analysis of single mechanosensory units confirmed no difference in firing frequency or activation threshold in response to distension between control and RTX bladder afferents (**Fig S6**), nor any differences in the firing patterns of low- and high-threshold subtypes (**Fig 3D-E**) including peak afferent response (**Fig 3Dii, 3Eii**), activation threshold (**Fig 3Diii, 3Eiii**), or area under the curve (AUC) (**Fig S7**). Aligning with the lack of effect of RTX on mechanosensory afferent responses or changes in bladder compliance, RTX had no impact on *in vivo* bladder function (**Fig 3G**). Quantification of *in vivo* bladder voiding behaviour found no difference in the number of void spots (**Fig 3Gi-iii**), and no differences in the proportion of small or large void spots (**Fig 3Gii-iii**) between control and RTX-treated mice. Together these findings indicate that mucosal afferents are not significant mediators of bladder distension-induced mechanosensory responses and bladder function under normal physiological conditions.

**Figure 3:**
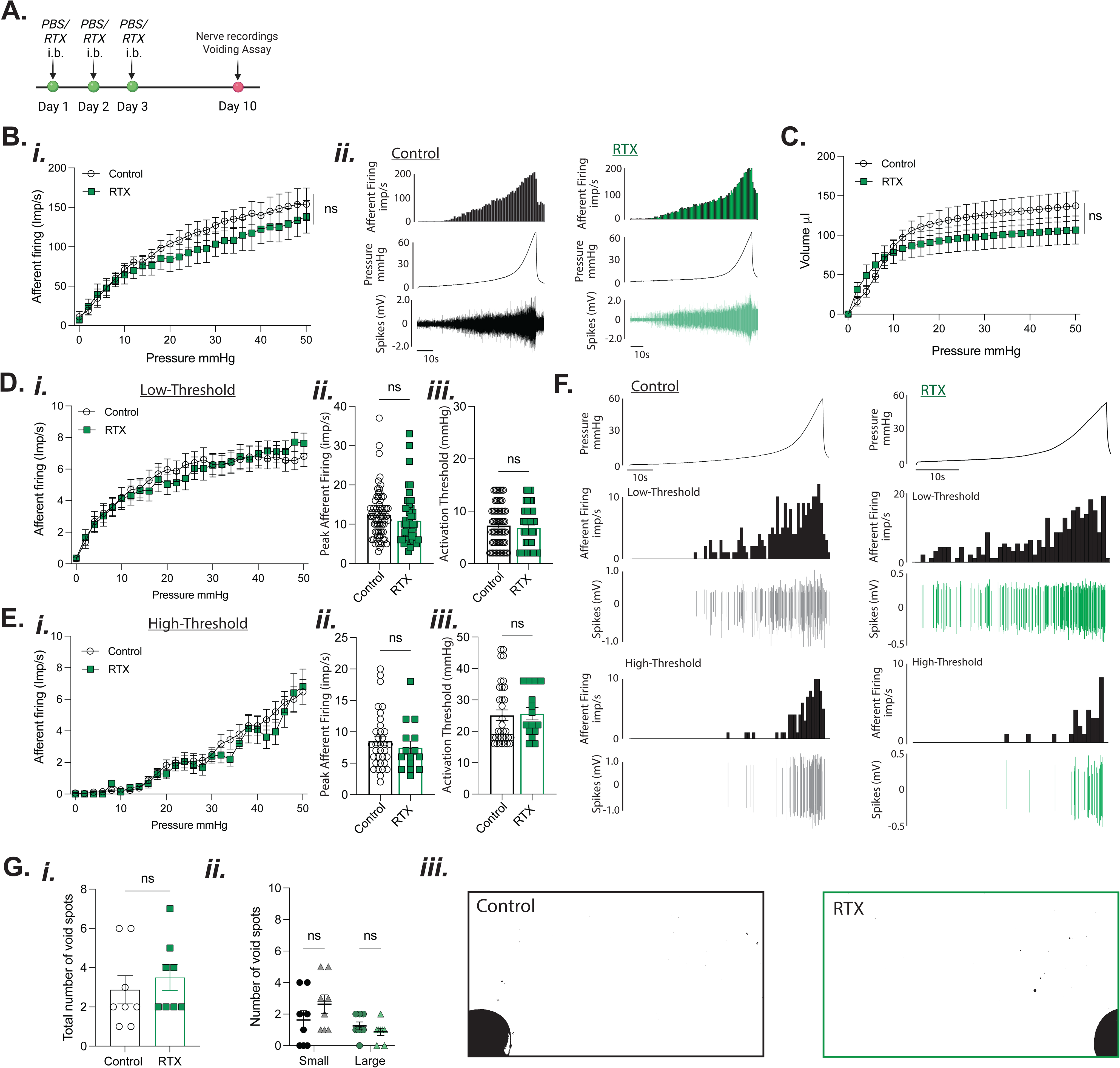
Mucosal afferent denervation does not impact bladder afferent mechanosensitive responses to distension *ex vivo* or *in vivo* bladder voiding patterns. **A)** Schematic representation of the experimental design. Mice received either RTX or PBS (SHAM) for 3 consecutive days and bladders were assessed 7 days after the final infusion. **(B)** *Ex-vivo* bladder afferent sensitivity to graded distension (0-50 mmHg) in control- and RTX-treated mice (**Bi)** (N=5). (**Bii**) Representative ex-vivo bladder afferent experimental trace showing intravesical pressure (mmHg), raw spike activity (mV), and afferent firing (imp/s) in response to graded bladder distensions in control (black) and RTX-treated (green) bladders. **(C)** Bladder muscle pressure/volume relationship (N=5). Single units were extracted from multiunit bladder afferent recording experiments and separated based on their activation threshold to distension as either low- **(D),** or high-threshold **(E)**. **(Di)** Low-threshold bladder afferent responses to distension; (**Eii**) maximum firing rate during distension; (**Diii**) and activation threshold (mmHg) (n= 52-69 individual units from N=5). (**Ei**) High-threshold bladder afferent responses to distension; (**Eii**) maximum firing rate during distension; (**Eiii**) and activation threshold (mmHg) (n= 15-34 individual units from N=5). **(F)** Representative experimental traces showing mechanosensitivity of low-threshold and high-threshold afferent units in response to graded bladder distensions (0-50 mmHg) from control (black) and RTX (green) treated bladders. (**G**) Analysis of voiding behaviour in control and RTX-treated mice including total void spots (**Gi)**, and number of small and large sized void spots **(Gii)** from control and RTX treated mice (N=8). **(Giii)** Representative images of voiding patterns in control and RTX treated mice. Data is presented as mean ± SEM. ^ns^ P > 0.05. Individual data points represent values from individual afferent units. Statistical differences for data in Ai, B, Ci, Di were determined by Two-way ANOVA with Šídák multiple comparisons. Statistical differences for data in Cii, Ciii, Dii, Diii were determined by unpaired Mann-Whitney test. Statistical differences for data in data in Fii were determined by two-way ANOVA with Bonferroni’s multiple comparisons.

### Denervation of mucosal afferents impacts bladder sensation and function during UTI and increases bacterial burden and inflammation

Despite their limited role in bladder function under healthy conditions, the dense sensory innervation of the mucosa (**Fig 1, Fig S1**) (9, 12) hints at a critical role in bladder homeostasis. Positioned near the bladder lumen, mucosal afferents are uniquely situated to rapidly detect infection or inflammation where it predominates during UTI (27, 28). Given that infection represents a significant evolutionary pressure, with early detection critical for host survival (29), we hypothesised that mucosal afferents may exist to detect pathological stimuli such as urinary tract infections (UTIs). To test this, we combined an established mouse model of UTI (1, 3) with our intravesical RTX model to selectively denervate mucosal afferents and evaluate their role in infection detection (**Fig 4A)**.

**Figure 4:**
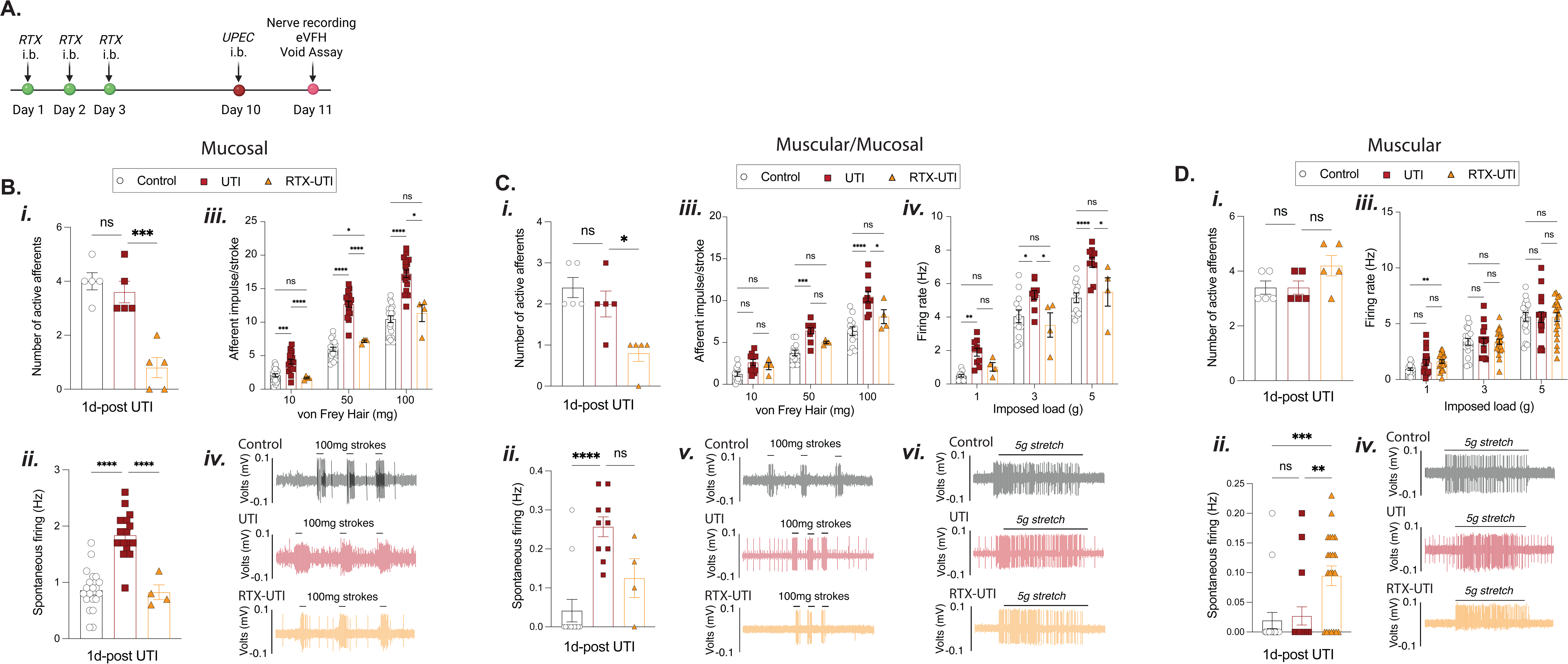
Mucosal but not muscular afferents are sensitised during UTI. (**A**) Schematic representation of the experimental design for the combined murine mucosal denervation and UTI infection model. **(B) Mucosal afferents**: (**Bi**) The number of mucosal afferents identified from individual experiments (control (black) n=20, UTI (red) n=18, RTX-UTI (orange) n=4, from N = 5 mice). (**Aii**) Spontaneous firing frequency of mucosal afferents. **(Biii)** Mucosal afferent responses to von Frey Hair stroking (10, 50, 100 mg). **(Aiv)** Representative mucosal afferent nerve recording traces from control (black), UTI (Red) and RTX-UTI (orange) treated bladders in response to 100 mg mucosal stroking. **(C) Muscular-Mucosal afferents**: **(Ci)** The number of mucosal nerves identified from individual experiments (control (black) n=12, UTI (red) n=10, RTX-UTI (orange) n=4, from N = 5 mice). (**Cii**) Spontaneous firing frequency of muscular-mucosal afferents. **(Ciii)** Muscular-mucosal afferent responses to von Frey Hair stroking (10, 50, 100 mg). **(Civ)** Muscular-mucosal afferent responses to bladder stretch by imposed load (1, 3, 5g). Representative bladder afferent nerve recording profiles of control (black), UTI (red) and RTX-UTI (orange) treated bladders to 100 mg mucosal stroking (**Cv**) and 5 g bladder stretch (**Cvi**). **(D) Muscular afferents: (Di**) The number of muscular nerves identified from individual experiments (control (black) n=17, UTI (red) n=17, RTX-UTI (orange) n=21, from N = 5 mice). (**Dii**) Spontaneous firing frequency of muscular afferents. **(Diii)** Muscular afferent responses to bladder stretch by imposed load (1, 3, 5g). **(Div)** Representative muscular afferent nerve recording traces from control (black), UTI (Red) and RTX-UTI (orange) treated bladders in response to 5g stretch. Data is presented as mean ± SEM. ^ns^ P > 0.05, * p < 0.05, **p < 0.01, ***p < 0.001, and ****p < 0.0001. Statistical differences for Ai, Bi, Ci were determined by Kruskal-Wallis test with Dunn’s multiple comparisons. For Aii, Bii, Cii, statistical differences were determined by Brown-Forsythe and Welch ANOVA test with Dunnett T3 multiple comparisons. For Aiii, Biii, Bv, Ciii statistical differences were determined by two-way ANOVA with Geisser-Greenhouse correction and Tukey’s multiple comparisons.

24hrs after bladder inoculation with uropathogenic *E. coli* (UPEC), mice developed acute UTI with bacteria detectable within the urine (**Fig S8**), a robust immune response characterised by neutrophil and inflammatory monocyte infiltration (16), and significant bladder damage and oedema that predominates in the urothelium and lamina propria (16, 27, 28) (**Fig S9**). *Ex vivo* bladder afferent nerve recordings revealed mucosal afferents exhibit significant increases in the frequency of spontaneous firing (**Fig 4Bii**) and significant hypersensitivity to mucosal stroking 24hrs after UPEC infusion compared to non-infected mice (**Fig 4Biii, iv**). Additionally, muscular-mucosal afferents exhibit increased spontaneous firing (**Fig 4Cii**) and develop significant hypersensitivity to both mucosal stroking (**Fig 4Ciii, Cv**) and stretch during UTI (**Fig 4Civ, Cvi**). In contrast, muscular afferents showed no differences in spontaneous firing (**Fig 4Dii**) or responses to bladder stretch following UPEC instillation (**Fig 4Diii, Div**). Intravesical RTX treatment prior to UPEC instillation (RTX-UTI) prevented UTI-induced sensitisation of mucosal (**Fig 4Bii-iv**) and muscular-mucosal afferent mechanosensory responses (**Fig 4Cii-vi**). Mucosal denervation prior to UTI induced a significant increase in spontaneous firing but had no impact on the mechanosensitivity of muscular afferents during UTI (**Fig 4D**). Together these data implicate mucosal afferents as major drivers of sensory signalling from the bladder during UTI.

In humans, UTI’s are typically associated with the development of distinct bladder sensations, including urinary urgency, dysuria, and pelvic pain which drive protective behaviours including an increase in the frequency of urination that can be recreated in mouse models (1, 2, 6–8, 18). To determine the consequences of inhibiting mucosal sensory signalling during UTI on the development of these protective behavioural responses, we assessed the development of pelvic pain and bladder dysfunction in UTI and RTX-UTI treated mice (**Fig 5**). Our data show that mice develop pelvic hyperalgesia (**Fig 5A**) and urinary frequency (**Fig 5Bi**), characterised by increased numbers of small urine spots during voiding analysis (**Fig 5Bii**) one day after UPEC instillation. Intravesical RTX treatment prior to UPEC instillation mitigates the development of pelvic hyperalgesia (**Fig 5A**) and prevents the development of urinary frequency (**Fig 5Bi, 5Bii, 5C**). Behavioural responses to UTI are considered important in infection clearance by provoking more rapid excretion of UPEC growing in the urine, along with exfoliated infected urothelial cells (16). To explore the consequences of inhibiting this host response on bacterial clearance, we compared the burden and extent of UTI following UPEC instillation in control UTI and RTX treated (RTX-UTI) mice (**Fig 5D**). UPEC was found to be increased within the bladder wall, urine, and kidneys (**Fig 5D)**, with a greater proportion (100% vs 40% in control UTI) of mice showing infection dissemination to the kidneys following mucosal denervation with RTX (**Fig 5Diii**).

**Figure 5:**
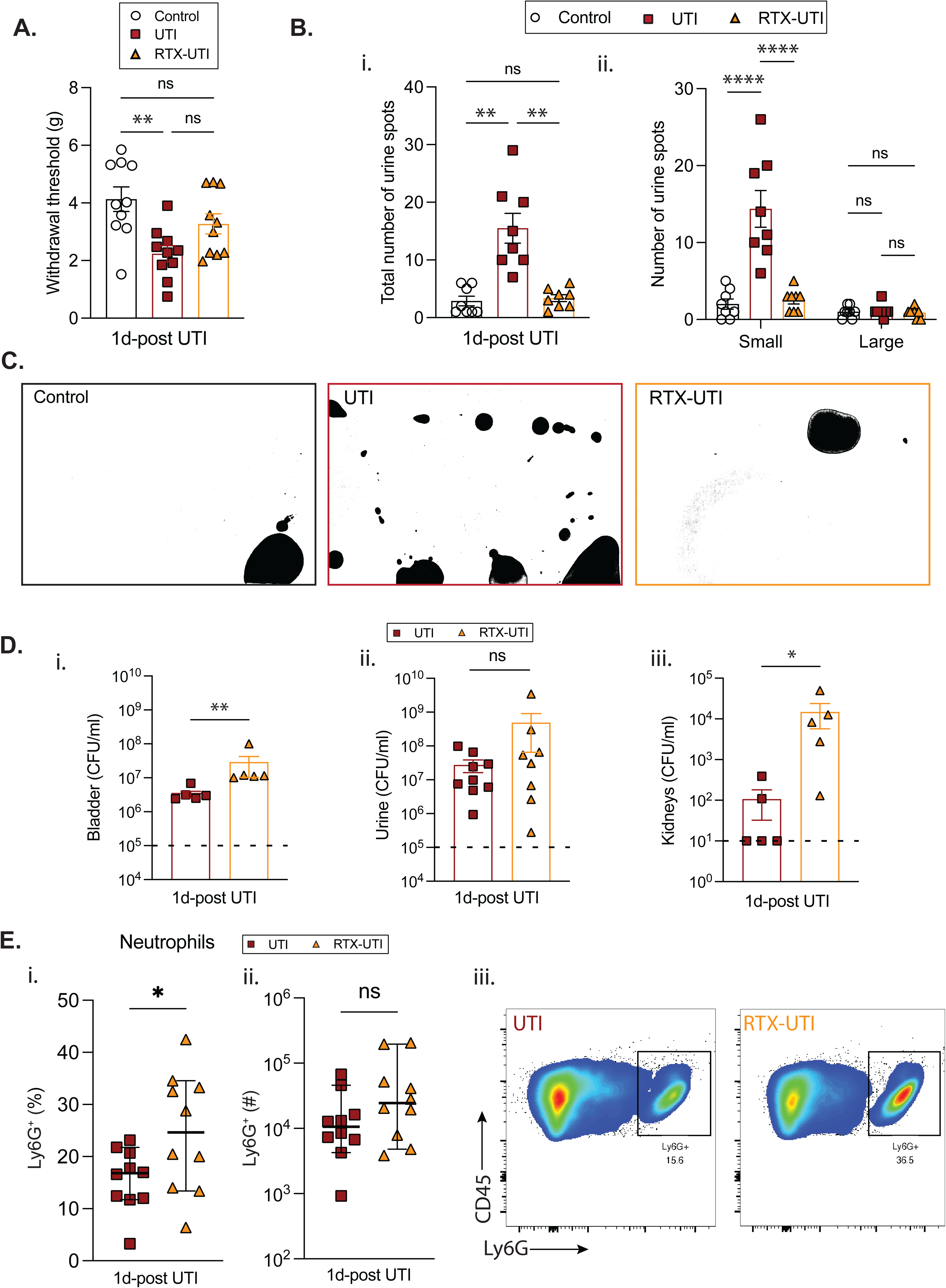
Mucosal afferents regulate bladder host-defence responses. (**A)** Withdrawal thresholds (g) to eVFH stimulation (N = 10). (**B**) Analysis of voiding behaviour including total void spots (**Bi)**, and number of small and large sized void spots **(Bii)** from control, RTX, and RTX-UTI treated mice (N=8). **(C)** Representative images of voiding patterns. **D**) Bacterial load in bladder (**Di**), urine (**Dii**), and kidneys (**Diii**) 24hrs after mice were infected with UPEC (N = 5-10). Dotted line represents the limit of detection for urine (10^5^), bladder (10^2^), kidneys (10^1^). **(E)** Neutrophil frequency as a percentage of total viable bladder immune (CD45^+^) cells (**Ei**) and total bladder neutrophil counts (**Eii**) in UTI and RTX-UTI mice. (**Eiii**) Representative flow cytometry plots of Ly6G^+^ neutrophils from UTI and RTX-UTI mice 24hrs after UPEC infection. Data is presented as mean ± SEM, ^ns^ P > 0.05, * p < 0.05, **p < 0.01, ****p < 0.0001. Individual data points represent values from a single mouse. Statistical differences for A, B were determined by Brown-Forsythe and Welch ANOVA tests with Dunnett T3 multiple comparisons. For D statistical differences were determined by unpaired t-test. Data in Ei analysed by Welch’s t-test and Eii by unpaired Mann-Whitney test.

The host-defence response to UTI includes a substantial innate immune response that is necessary for infection clearance (16, 27, 28). To assess whether mucosal denervation with RTX changed this response we utilised flow cytometry to assess the frequency of major immune cell populations including T cells, monocytes, macrophages, neutrophils and NK Cells (**Fig 5, Fig S10**). The increase in UPEC burden seen in the bladders of RTX-UTI mice is associated with a significant increase in the frequency of neutrophils in the bladder wall (**Fig 5Ei,iii)** without significant changes in the frequency or number of other immune cell populations **(Fig S10**).

## Discussion

In health, bladder sensation primarily reflects the degree of bladder fullness, gradually intensifying as the bladder fills until voiding is initiated (7, 8). During a urinary tract infection (UTI), however, the relationship between bladder volume and sensation becomes dysregulated, with bladder sensations dominated by irritative and painful sensations, including pelvic pain/pressure, dysuria, and urgency even when the bladder is not full (8, 33). In this study, we present data showing that discrete functional subclasses of bladder afferents differentially contribute to sensation and bladder function in health versus disease. Using a novel mouse model of mucosal denervation, we demonstrate that muscular afferents govern bladder mechanosensation and voiding function in health while mucosal afferents emerge as the dominant drivers of sensory signalling during infection to initiate urinary frequency that contributes to the host-defence response to infection.

As normal function of the bladder is related to the accurate detection of bladder muscle stretch as a proxy for bladder fullness, the role of stretch-insensitive mucosal sensory nerves in bladder sensation and function has been an enduring question since their discovery almost 20 years ago (4, 5). Experimental interrogation of mucosal afferents has been constrained by two major limitations: first, there are no molecular markers that reliably distinguish muscular and mucosal afferents, precluding their selective chemogenetic or cre-inducible ablation (23, 30, 31); and second, the absence of clear anatomical segregation by spinal level (5, 6) blunts the potential of recently developed techniques for targeted dorsal root ganglia (DRG) removal (32). RTX has long been used as a molecular scalpel to ablate TRPV1⁺ nerves via central or systemic administration (21, 23). However, because TRPV1 is widely expressed throughout bladder afferents (24, 25, 33), these approaches lack the spatial precision needed to study mucosal afferent function in isolation. To overcome this limitation, we developed a novel model of spatially selective mucosal denervation by infusing RTX intravesically into the bladder over 3 consecutive days. This approach produced a robust and specific attenuation of mucosal afferent signalling, and reductions in CGRP⁺ and TdTom⁺ mucosal innervation whilst preserving muscular afferent innervation and function. This model thus enabled a systematic dissection of the functional specialisation of muscular and mucosal nerves. Using graded bladder distension as a proxy for normal bladder filling, and spontaneous voiding as an assessment of healthy bladder function, we found pelvic afferent responses and bladder function in healthy control mice were unchanged following mucosal denervation. In contrast, we show mucosal afferents are crucial for signalling the presence of UTI, inducing referred pelvic pain, and increasing urinary frequency.

Sensory nerves have a well-known role in detecting and signalling pathological stimuli from both the skin and viscera to elicit unique sensations and protective behavioural responses (29). Infection and inflammation are classical drivers of neuronal sensitisation (26, 34), and both bacterial and inflammatory stimuli have been shown to sensitise bladder DRG (2, 3) and induce pelvic hyperalgesia and urinary frequency (1, 2, 18, 35). How mucosal innervating afferents are exclusively sensitised during UTI, however, remains unclear. It is possible that mucosal afferents exhibit a unique phenotype that mediates responses to stimuli of distinct modalities through unique ion-channel/receptor profiles in line with the specificity theory of labelled lines (36). In support, muscular and mucosal afferents in mice have known differences in mechanosensitivity (4, 5), neuropeptide content (12, 37), and sensitivity to chemical stimuli (13–15) that hint at unique molecular profiles that may underpin a unique function. An alternative proposition is that the functional specificity of mucosal afferents we observed may be directly related to their location within the urothelium and lamina propria, where infection and inflammation predominate during UTI (16). Patients with clinically distinct forms of non-infectious cystitis, including interstitial, radio-, immuno-, and chemotherapy-induced cystitis typically describe bladder sensations and symptoms in line with those experienced during UTI (38–40), suggesting the same neural mechanisms may underlie these sensations and implicating mucosal afferents as broad sensors of pathological stimuli, including both infection and inflammation. Similarly, mucosal afferents are preferentially sensitised in a non-infectious guinea pig cystitis model, and cystitis induced nocifensive visceromotor responses to bladder distension are reduced following acute intravesical instillation of RTX (41). Mucosal afferents can also be sensitised or respond directly to a broad variety of inflammatory molecules (13–15); and undergo immune-mediated sprouting that drives pelvic sensitivity and urinary frequency after recurrent UTI (35). When spatial proximity is removed, such as with isolated non-specific populations of bladder innervating DRG, we see non-specific sensitisation by supernatants from infected bladders (2). Clinically, studies in patients confirm that specific targeting of sensory nerves via the bladder lumen with phenazopyridine, lidocaine, and RTX preferentially and effectively relieve hypersensitivity symptoms including urgency and bladder pain without major impacts on normal bladder function (42–47), further supporting a major and specific role for mucosal afferents in bladder pain signalling.

Irritative sensations driven by the activation of peripheral sensory nerves are crucial signifiers of infection and inflammation that are necessary for directing behavioural responses to protect against threats across various organs (48, 49). Urinary frequency has been proposed to act as an analogous defensive behavioural response that aids the innate immune response in bacterial clearance through rapid expulsion of bacteria that would otherwise be stored and growing in the bladder reservoir (19, 20). Clinically, disruptions to bladder sensory or motor functions such as those observed in persons with neurogenic lower urinary tract dysfunction caused by Parkinson’s disease, multiple sclerosis, spinal cord injury, and diabetic neuropathy are associated with increased susceptibility to both initial and recurrent UTI’s, and an increased risk developing severe UTI’s (50–54). In this study, we show that mucosal afferents are essential for driving the development of urinary frequency during UTI. Furthermore, in the absence of UTI-induced urinary frequency following mucosal denervation, we observed greater bacterial burden in the bladder wall and kidneys, providing evidence of a mechanistic link between altered bladder sensation and UTI burden.

In addition to the direct detection of threats through the generation of sensations that drive protective behaviours, sensory nerves have also been implicated in host-defence responses in a variety of organs via neuro-immune interactions driven by local antidromic release of neuropeptides, including CGRP and Substance P from afferent endings (55). In this study, RTX caused an almost complete loss of the neuropeptide CGRP in the mucosa, however, aside from an increase in the frequency of neutrophils, no other changes in immune cell infiltration were observed during UTI between RTX and sham treated groups. CGRP has been shown to directly act on immune cells to inhibit cytokine release and modulate neutrophil migration (31, 55, 56). Neutrophils are also the primary effector cells responsible for bacterial clearance, and their infiltration typically correlates with infection severity (57). Consistent with this, we observed that increased neutrophil frequency was associated with greater bacterial burden in the bladder, aligning with a direct increase in the immune response to counter an increased severity of infection (57, 58). Whether bladder neuropeptides also directly regulate immune responses during UTI will require future studies focused exclusively on neuropeptide depletion without compounding mucosal denervation.

Together, our data implicate bladder mucosal afferents as key sensors of pathogenic challenge and reveal a previously unrecognised mechanism that drives behavioural responses to reduce the burden and extent of UTI.

## Materials and Methods

### Animals

The Animal Ethics Committees (AECs) of the South Australian Health and Medical Research Institute (SAHMRI) and Flinders University approved all procedures (Approvals #21-038; #5844, #6887). Female mice aged 6-8 weeks of age were used for all experiments and acquired from in-house C57BL/6J breeding programmes within SAHMRI’s specific and opportunistic pathogen-free animal care facility and College of Medicine and Public Health Animal Facility (COMPHAF). Mice were housed (up to 5 mice per cage) in a temperature-controlled environment of 22°C and a 12-hour light/12-hour dark cycle within ventilated cages with free access to food and water. Mice were randomly assigned to experimental procedures and were single housed where necessary for experimental design.

Na_v_1.8^-Cre-TdTomato^ mice were created as previously described (59) by crossing Na_v_1.8^-Cre^ mice with TdTomato reporter floxed mice. Transgenic mice on a C57Bl/6 background expressing Cre recombinase under control of the Na_V_1.8 promoter were generously donated by A/Prof Wendy Imlach (Monash University, Melbourne, Australia) and were originally generated from JAX stock #036564 (60). Na_V_1.8-Cre mice were crossed with tdTomato reporter mice gifted by Prof Susan Woods (SAHMRI, Adelaide, Australia); these mice possess a loxP-flanked Stop cassette that is excised in Cre-expressing neurons, allowing tdTomato production in cells expressing Na_V_1.8-Cre only. As it has been previously shown that Na_v_1.8 is present in the vast majority of bladder neurons (61), this mouse provides an effective and specific sensory neuron marker in the bladder.

### Resiniferatoxin treatments

Resiniferatoxin (RTX; 1mg) was dissolved in dimethyl sulfoxide (DMSO) to a stock concentration of 1mM. 50µl of frozen stock was diluted in 1.5ml of phosphate buffered saline (PBS) to give a 30µM working concentration to perform intravesical RTX infusions in mice. 6-8 week old female C57BL/6J or Na_v_1.8^-Cre-TdTomato^ mice were anaesthetised under isoflurane (3%/1L O_2_) and a lubricated catheter was inserted into the bladder via the urethra. Urine was drained from the bladder followed by instillation of 100µl of 30µM RTX for 30 mins. Control mice received sham treatments consisting of bladder catheterisation and infusion of 100µl of 1X PBS. Mice received the same infusions across three consecutive days and utilised for downstream experiments 7 days after the final infusion.

### Quantification of bladder nerve density

Bladders were surgically removed from RTX and control mice and fixed in 4% PFA (Sigma-Aldrich, #158127) for 18 to 20 hrs at 4°C. PFA was removed, and the tissue was cryoprotected in 30% sucrose/phosphate buffer for 18 to 20 hrs at 4°C prior to freezing in 100% Optimum Cutting Temperature (OCT) Compound (Sakura Finetek) using liquid nitrogen cooled 2-methylbutane (Sigma-Aldrich, M32631). Bladders were frozen in intermediate sized cryomolds (Tissue-Tek, Sakura Finetek) with the bladder dome orientated towards the top of the cryomold. Bladders were cryosectioned at 40µm thickness and placed on gelatine-coated slides for immunofluorescence labelling. Bladders were serially sectioned from the urethral opening to the dome over 4 slides.

### Calcitonin Gene Related Peptide (CGRP) Immunohistochemistry

Bladder sections from Na_v_1.8^-Cre-tdTomato^ mice were air dried for 20 mins prior to three washes with 0.2% Triton X-100 (Sigma-Aldrich, cat. T8787) in 1X PBS (PBSTx_0.2%_) for 5 mins each. Sections were incubated in 5% normal chicken serum (Thermo Fisher Scientific-Gibco™, cat # C34777) diluted in PBSTx_0.2%_ for 30 mins at room temperature and then incubated with primary CGRP antibody (1:400) (Sigma-Aldrich, cat #PC205L) at room temperature for 12 to 18 hours in a moist, wet and humid environment. Sections were washed three times with PBSTx_0.2%_ for 5 mins each prior to incubation for 1 hr with chicken anti-rabbit-AF488 secondary antibody (1:200) (Thermo Fisher Scientific-Invitrogen Cat #A-21441) at room temperature. Following three 5 min washes with PBSTx_0.2%_, sections were cover slipped with Prolong Diamond Antifade Mountant with DAPI (Thermo Fisher Scientific-Invitrogen; cat. P36962). Slides were left to dry at room temperature for 24 hrs before visualisation. Bladder sections were visualised using the Zeiss Axio Scan.Z1 slide scanner at Adelaide Microscopy. Images were obtained with 1-4 manually set focus points per section at 20x magnification using the 475nm and 567nm LED module. To visualise Alexa Fluor 488 fluorescence, the 475nm module was tuned to 493nm excitation and 517nm emission settings. While TdTomato was visualised using the 567nm module tuned to 553nm excitation and 568nm emission settings. These settings were saved and used for all imaging. Images were automatically saved as a Carl Zeiss Image and viewed using Zen Blue. Images of sections from each region (opening, middle and dome) were exported as a TIFF file.

### Analysis

Images were analysed in ImageJ using the area tool. The Integrated Density (IntDen), area and %area was measured for the mucosal and detrusor layer for each mouse and were based on the TdTomato and AF488 fluorescence that was above a set threshold. The IntDen is the product of the area and mean gray value, the area is the area of the selection in mm^2^ and %area is the percentage of pixels in the image or selection that is above a set threshold. The IntDen, area and %area measurements were averaged across each bladder region (opening, middle and dome). Consistency of area taken for analysis across groups was confirmed by assessing perimeter of mucosa and detrusor using ImageJ (**Fig S11**).

### CGRP ELISA

Bladders were removed from RTX and SHAM treated mice and mucosa and detrusor were rapidly separated by fine dissection and snap frozen in liquid nitrogen. Frozen tissues were weighed and 10µl 100% acetic acid/mg tissue was added before bead homogenised using the TissueLyserII (Qiagen, SN: 125090912) for 6 min at 30Hz. Samples were briefly centrifuged and stored at 4°C for 15 to 20 mins. Homogenates were ultracentrifuged (45000 rpm for 30 mins at 4°C) and the supernatant was carefully removed without disturbing the pellet and transferred to a clean Eppendorf tube. Bladder protein extracts were stored at -20°C until future use.

Bladder protein extracts in supernatants were quantified using the Invitrogen EZQ™ Protein Quantitation Kit (Thermo Fisher Scientific, cat. R33200) according to manufacturer’s instructions. The CGRP (rat) ELISA kit (Bertin Bioreagent, cat. A05482) was used to quantify the amount of CGRP within the supernatants as per manufacturer’s instructions. All bladder protein extracts were diluted to 20 mg/ml using the kit’s EIA buffer to standardise the concentration of protein when loading the ELISA plate. 100ml of samples, standards, and controls (blank, non-specific binding wells, quality control wells) were plated as duplicates. The plate was incubated for 16 hrs at 4°C, rinsed with wash buffer three times, and blotted dry before Ellman’s reagent was added to each well. The plate was then incubated in the dark at room temperature on an orbital shaker at 50 rpm before being read at 405 nm after 1 hr (Beckman DTX 880 Plate Reader (Beckman Coulter).

### Flow cytometry

Bladders were surgically removed and vertically bisected along the urethra towards the bladder dome and placed in an Eppendorf tube containing 1ml of RPBMI1640 medium containing digest buffer (1mg/ml collagenase + 100µg/ml DNAse I) (Thermo Fisher Scientific™ - Gibco™, cat. 17100017; Bio-Rad, cat. 7326828). Bladders were cut into small pieces with dissection scissors and then placed into a shaking incubator for 1 hr at 37°C at 180 rpm. Digestion was stopped by adding 1ml of cold RPMI (Thermo Fisher Scientific™ - Gibco™, cat. 11875093) with 10% Fetal Bovine Serum (FBS) (Thermo Fisher Scientific™ - Gibco™, cat. 10100147). The tissue suspension was then pushed through a 70 µm cell strainer and then centrifuged at 400 x g for 5 mins to pellet the cells. The supernatant was then aspirated, and the cell pellet was resuspended in 800µl of BD Pharm Lyse red cell lysis buffer (Becton Dickinson (BD) Biosciences, cat. 555899) at room temperature for 90s before 1ml of RPMI with 10% FCS was added to prevent further cell lysis. Cells were centrifuged at 400 x g for 5min, and the cell pellet was resuspended in 200µl of FACS buffer (0.1% Bovine Serum Albumin (BSA) and 0.4% Ethylenediaminetetraacetic Acid (EDTA) in PBS). Resuspended cells were transferred to a 96-well round-bottom plate, with samples from each bladder going into one well, and centrifuged at 300 x g for 5 min. The supernatant was removed by gently tapping the plate before being resuspended in 50µl of a previously prepared primary antibody cocktail (Table 1) and left to incubate for 30 mins on ice in the dark. Cells were rinsed with 200µl of FACS buffer and centrifuged at 300 x g for 5 mins at 4°C. Wells were then resuspended in 50µl of secondary antibody (Steptavidin-PE-CSF594; BD Biosciences) diluted 1:500 in FACS buffer; the cells were left to incubate with the secondary antibody for 15 to 20 mins on ice in the dark. After which the cells were washed twice with 200µl of FACs buffer and then resuspended in 120µl of FACs buffer in preparation for flow cytometry analysis. Lastly, 10µl of DAPI (Sigma-Aldrich, cat. 124653) and 4µl of counting beads were added to each well. Flow cytometry was performed using the BD LSRFortessaTM X-20 (BD Bioscience) at the SAHMRI flow cytometry core facility using the gating strategy shown in **Figure S12**. Subsequent data analysis was conducted using FlowJo software version 10.8.1.

**Table 1:** Pan Immune Cell Bladder Antibody Panel.

| Target | Fluorophore | Company | Cat# | Dilution | Marker/function |
| --- | --- | --- | --- | --- | --- |
| **Fc Block** | - | BD Biosciences | 553141 | 1:400 | Used to block non-specific binding of antibodies to cells |
| **CD45.2** | APC-Cy7 | BD Biosciences | 560694 | 1:400 | Expressed on many different types of immune cells |
| **CD19** | PerCP-Cy5.5 | BD Biosciences | 551001 | 1:200 | Specific to B cells |
| **CD3e** | PerCP-Cy5.5 | BD Biosciences | 551163 | 1:200 | Present on T cells |
| **CD4** | BV510 | BD Biosciences | 563106 | 1:200 | Found on CD4 T cells, a subtype of helper T cells |
| **CD8a** | BUV395 | BD Biosciences | 563786 | 1:200 | Found on CD8 T cells, a subtype of cytotoxic T cells |
| **NK-1.1** | APC | BD Biosciences | 550627 | 1:200 | A specific marker for Natural Killer (NK) cells |
| **Ly-6G** | Pe-Cy7 | BD Biosciences | 560601 | 1:400 | Specific marker for neutrophils |
| **CD11b** | PE | BD Biosciences | 557397 | 1:800 | Specific to myeloid cells |
| **CD11c** | Biotin | Miltenyi | 130-101-929 | 1:50 | Expressed on monocytes, granulocytes, and tissue macrophages |
| **MHCII (I-A/I-E)** | BV711 | BD Biosciences | 563414 | 1:1500 | Found on macrophages, B cells, and other myeloid cells; indicates activation in myeloid cells |
| **Ly-6C** | FITC | BD Biosciences | 553104 | 1:400 | Specific to monocytes |
| **F4/80** | BV421 | BD Biosciences | 565411 | 1:400 | Specific to macrophages |

### Haematoxylin and Eosin staining

Bladders were collected and fixed in 4% PFA for 24 hrs, rinsed, and transferred to 70% ethanol. Bladders were then embedded in paraffin and serially sectioned from the middle of the bladder towards the dome in 5µm sections and subsequently stained with H&E by the Flinders Microscopy and Microanalysis Core Facility.

Bladder H&E-stained sections were visualised using a slide scanner (Olympus VS200, Olympus Corporation, Tokyo, Japan) at Flinders Microscopy and Microanalysis Core Facility. Images were obtained with 1-4 manually set focus points per section at 20x magnification using brightfield imaging settings. Sections were scored to compare inflammation between the bladder mucosa and muscle by a researcher blinded to the conditions after training by a pathologist (Prof Sonja Klebe). Sections were scored according to Table 2.

**Table 2:** Histopathological score criteria for bladder muscle and mucosa.

**Table : Histopathological score criteria for bladder muscle and mucosa**
| Score | Tissue damage |  | Oedema |  | Haemorrhage |
| --- | --- | --- | --- | --- | --- |
|  | Muscle (fibrosis) | Mucosa (sloughing) | Muscle | Mucosa |  |
| 0 | Full muscle integrity | Full urothelium integrity | No oedema present | No oedema present | No free erythrocytes |
| 1 | Minimal fibrosis (<25% disruption of muscle fibres) | Sloughed urothelium (<25% of section) | Slight separation of muscle fibres | Slight separation in urothelial cells | <10 erythrocytes |
| 2 | Moderate fibrosis (25-50% disruption) | Sloughed urothelium (25-50% of section) | Moderate separation of muscle fibres | Moderate separation, expanded submucosa | 10-30 erythrocytes |
| 3 | Severe fibrosis (>50% of muscle fibre disruption) | Sloughed urothelium (>50% of section) | Severe stretching/distortion of muscle fibres | Severe separation, stretching/distortion of urothelial cells | >30 erythrocytes |

### *Ex vivo* bladder pelvic afferent nerve recordings

#### Flat-sheet bladder preparation

To individually evaluate the direct activity of the bladder mucosal and muscular nerves, *ex vivo* flat-sheet bladder afferent recordings were performed (41, 62, 63). After mice were humanely killed via CO_2_ inhalation, the bladder and associated connective tissue containing nerve bundles was dissected out and opened along the midline of the anterior wall from the urethra up to the apex into a flat sheet in Krebs buffer solution (composition in mmol/L: 118.4 NaCl, 24.9 NaHCO_3_, 1.9 CaCl_2_, 1.2 MgSO_4_, 4.7 KCl, 1.2 KH_2_PO_4_, and 11.7 glucose) bubbled in 95% oxygen and in 5% carbon dioxide. Nerve trunks entering the trigone area of the bladder between the left (or right) ureter and urethra were carefully isolated from surrounding connective tissue using a tungsten needle. The bladder was then orientated into a rectangular shape (approximately 12mm wide by 15mm long), encompassing the bladder trigone and body and pinned mucosal side up with the nerves along one edge in a 22ml organ bath continuously perfused with warmed 34°C Krebs (3ml/min). The nerve trunks were pinned loosely with 50µm gold-plated tungsten pins, and a ring of paraffin was placed around and over the nerves; this formed the foundation for making a paraffin oil bubble for electrical isolation and recording of electrical activity. The opposite edge of the bladder was attached via a hook and cantilever system, to an isotonic transducer (Harvard Bioscience 52-9511, S Natick, MA, USA) for stretching imposed by loads (1-5g) and simultaneous measuring of the preparation lengthening. The preparation was rested for 30 min before commencing experiments. Electrical signals were amplified (DAM 80, WPI), filtered via band-pass filter (BPF-932, CWE, USA, band pass 10Hz-10kHz) and recorded by a computer at 20kHz with a Micro 1401-4 data acquisition system (CED, UK). Single units were discriminated offline by using Spike 2 version 10 software (CED, UK).

#### Characterisation of bladder afferent subtypes

The current study focussed on the three major functional classes of bladder afferents: mucosal, muscular-mucosal, and muscular afferents. Afferent sensitivity to stroking was determined using calibrated von Frey hairs (10, 50 and 100 mg) stroked across the receptive field at a rate of 5mm/s 5 times. The middle three strokes were used for analysis. Afferent sensitivity to bladder stretch was determined by imposed loads (1, 3 and 5g) applied via the cantilever system for 10s with a 1 min interval between each load. Afferent subtypes were differentiated based on their well characterised responsiveness to stroking and stretching (13, 15). Mucosal afferents respond to light mucosal stroking with von Frey Hairs but not to bladder stretch imposed by a load. Muscular afferents do not respond to von Frey hair stroking but do respond to bladder stretch. Muscular-mucosal afferents respond to both mucosal stroking by von Frey hairs and to bladder stretch.

#### Analysis of bladder afferent activity

Nerve recording activity was analysed offline using Spike2 software. Single units were first discriminated using defined parameters in Spike2. To analyse spontaneous activity or stretch the ‘rate’ channel mode was selected with 0.5 seconds time width, while for stroking activity the ‘lines’ channel mode was selected. The total number of units within a stroking or stretching event was calculated.

### Whole bladder pelvic afferent recordings

To evaluate the nerve activity of the bladder during distension, *ex vivo* whole bladder afferent recordings were used as previously described (24, 33, 59, 61, 64–66). Mice were humanely killed by CO_2_ inhalation, and the entire lower abdomen was removed and submerged in a modified organ bath under continual perfusion with gassed (95% O_2_ and 5% CO_2_) Krebs buffer at 35°C. The bladder, urethra, and ureters were exposed by removing excess tissue. Ureters were tied with 4-0 perma-hand silk (Ethicon; cat. LA53G). The bladder was catheterised (PE 50 tubing) through the urethra and connected to a syringe pump (NE-1000) to allow a controlled fill rate of 100µl/minute with saline (NaCl, 0.9%). A second catheter was inserted through the dome of the bladder, secured with silk, and connected to a pressure transducer (Digitimer; cat. NL108T2) to enable intravesical pressure recording during graded distension. Pelvic nerves, isolated from all other nerve fibres between the pelvic ganglia and the spinal cord, were dissected into fine multiunit branches, and a single branch was placed within a sealed glass pipette containing a microelectrode (WPI) attached to a Neurolog headstage (Digitimer; cat. NL100AK). Nerve activity was amplified (NL104), filtered (NL 125/126, bandpass 50-5000 Hz, Neurolog; Digitimer), and digitised (Cambridge Electronic Design; cat. CED 1401) to a PC for offline analysis using Spike2 software. The number of action potentials crossing a preset threshold at twice the background electrical noise was determined per second to quantify the afferent response. Single-unit analysis was performed offline by matching individual spike waveforms through linear interpolation using Spike2 version 5.18 software. Afferent units were deemed ‘low threshold’ if continuous action potential firing was elicited at less than 16mmHg. By contrast, ‘high-threshold’ afferents displayed continuous action potential firing only when pressures exceed 16mmHg.

#### Afferent recording experimental protocols

At the start of each afferent recording experiment, control bladder distensions were performed with intravesical infusion of saline at a rate of 100µl/min to a maximum pressure of 50 mmHg at 10 min intervals to assess the viability of the preparation and reproducibility of the intravesical pressure and neuronal responses to distension. The volume in the bladder was extrapolated from the known fill rate (100µl/min) and the time taken (s) to reach maximum pressure (50 mmHg). Compliance was determined by plotting intravesical pressure against the calculated volume.

#### Analysis of bladder afferent activity

Nerve recording activity was analysed offline using Spike2 software. Afferent activity from multiunit nerve recordings was determined using established scripts as the ‘rate’ of action potential firing above the baseline threshold, which was set at twice the background electrical noise. Single units were first discriminated using defined parameters in Spike2 that were consistent across all experiments. Afferent activity from single units was assessed as ‘rate’ of firing per unit pressure

### Voiding paper assay

Voiding paper assay is a micturition assessment tool that provides information about changes in urinary behaviour and was performed as previously described (67–69). Single housed mice were temporarily removed from their cages and their bedding was replaced with filter paper (Whatman; Grade 1) fixed to the bottom of the cage. Mice remained in their lined cages for 3 hrs, between 10AM and 1PM, with free access to food and water after which filter paper was collected, imaged using an ultraviolet transilluminator (Gel Doc XR+ Gel Documentation System; Bio-Rad), and saved as .jpg files. Images were digitised into binary images using ImageJ software and the number and size of urine spots was determined using pre-set thresholds within the ImageJ software to identify all urine spots and exclude background. For the purposes of this study, the total number of urine spots per filter paper were quantified and the number of small (500 to 100,000 pixels) and large (>100000 pixels) spots were stratified.

### Electronic von Frey Hair (EvFH) testing

The electronic von Frey Hair (EvFH) test was used to assess abdominal sensitivity to mechanical stimuli by measuring the withdrawal response to a stimulus (67, 70). The BIO-EVF4 (Bioseb) was used to conduct EvFH tests. Prior to beginning EvFH tests, mice were habituated to the EvFH enclosure and placed on an elevated wire rack, for 30 mins each day, across 2 days. On the day of testing, mice were moved to the testing room and allowed to acclimatise for a minimum of 15 mins. Animals were individually placed on the wire rack within the enclosures and left undisturbed for a minimum of 15 mins. The EvFH unit was fitted with a semi-flexible tip and zeroed. The force threshold, the minimum force required for the instrument to commence recording, was established as 0.70g within the BIO-CIS software. Once the animal was still and quiet, the force transducer was applied perpendicularly to the animal’s lower abdomen from below. Force was gradually and linearly increased until a clear withdrawal response was recorded. The minimum force applied (in grams) that elicited the abdominal withdrawal was noted as the withdrawal threshold. The pelvic area was stimulated three times over 5 minutes, and an average withdrawal force was calculated from the trials. The mice were tested on rotation, so each mouse was stimulated first before starting the second round of stimulations to reduce acclimation to the filament.

### Mouse model of Urinary Tract Infection

#### UPEC culture and infusions

UPEC culture and infusions were performed as previously described (1). UPEC were cultured in sterile Luria-Bertani (LB) broth. UPEC broth was prepared using frozen UPEC stock (ATCC; CFT073 strain) a commonly used strain to establish UTI animal models (71–73). Single colonies were isolated following streak plate with sterile 10µl inoculation loop. The plate was cultured overnight in an incubator at 37°C. Single colonies were picked and grown in LB broth overnight in an orbital shaking incubator at 37°C at 200 rpm. A spectrophotometer (Perkin Elmer VICTOR X5 plate reader) was used to obtain an optical density (OD) at 600nm, of 0.20 ± 0.01 to give a 1 x 10^8^ CFU/ml concentration of bacteria.

UPEC instillation was performed in C57BL/6J mice aged 7-8 weeks old, 7 days after the final intravesical infusion of RTX (RTX-UTI), or PBS (UTI). Mice were anaesthetised with isoflurane and a lubricated sterile catheter was inserted into the bladder via the urethra and urine was removed. A second sterile catheter was used to instil 50µl of UPEC broth (1 x 10^8^ CFU/ml) for 10 mins. After 10 mins, the catheter was gently removed from the urethra leaving the UPEC broth in the bladder. Sham (control) mice received instillations of 50µl 1X PBS for 10 mins. 1 day after bladder infections, urine was obtained from mice via scruffing and collection of voided urine into a sterile Eppendorf. Mice were subsequently humanely killed via CO_2_ inhalation and bladders and kidneys were removed from mice and processed for assessing bacterial load.

Bladders and kidneys from each mouse were placed in FastPrep® microfuge lysing matrix tubes (MP Biomedicals, cat. 115076200-CF) containing ice cold sterile 1X PBS and a sterile chrome steel bead and lysed using The FastPrep-24™ (MP Biomedicals) until the tissues were adequately homogenised (40-80s at 4.0m/s) before being plated on agar. 100µl of the neat bladder and kidney homogenate and urine were plated on MacConkey No.3 Agar plates (Oxoid™; cat. CM0115B) prepared according to manufacturer’s instructions. Urine was first diluted 1:100 in sterile LB broth and then serially diluted at the 10^1^, 10^2^ and 10^3^ dilution factors and 100µl of each dilution was pipetted onto the corresponding MacConkey No.3 Agar plate and spread with a sterile L-shaped spreader until it was absorbed into the agar. This was replicated for all duplicates for all mice at each time point. The plates were then incubated at 37°C for 24 hrs before colony counting to assess bacterial load.

#### Statistical Analysis

GraphPad Prism v10.2.3 (GraphPad Software, La Jolla, California, USA) was used for statistical analyses, using Student’s t-tests when comparing two independent variables, or analysis of variance (ANOVA) when comparing three or more independent values. Normality tests were performed to determine whether a parametric or nonparametric statistical test should be used. If the data was normal a parametric test was used and if not normal, a nonparametric test was used. The significance value was set at 0.05.

## Acknowledgements

The authors acknowledge the facilities, and the scientific and technical assistance of Microscopy Australia (ROR: 042mm0k03) enabled by NCRIS and the government of South Australia at Flinders Microscopy and Microanalysis (ROR: 04z91ja70), Flinders University (ROR: 01kpzv902). We would like to thank SM Brierley for providing access to Nav1.8-TdTomato mice from their established breeding colony.

## Authorship contribution statement

**Cindy Tay**: Writing – review & editing, Writing – original draft, Visualization, Methodology, Investigation, Formal analysis, Conceptualization

**Harman Sharma**: Writing – review & editing, Writing – original draft, Visualization, Methodology, Investigation, Formal analysis, Conceptualization. **Stewart Ramsay:** Writing – review & editing, Writing – original draft, Visualization, Methodology, Investigation, Formal analysis. Georgia Bourlotos: Writing – review & editing, Investigation. Sarah K. Manning: Writing – review & editing, Investigation. Natalie E. Stevens: Writing – review & editing, Methodology, Investigation, Formal analysis. Feargal J. Ryan: Writing – review & editing, Methodology, Formal analysis, Conceptualization. Geraint B. Rogers: Supervision, Resources. David J. Lynn: Writing – review & editing, Supervision, Resources, Methodology. **Andrea M. Harrington:** Writing – review & editing, Supervision, Resources, Methodology, Investigation, Formal analysis, Conceptualization. **Vladimir Zagorodnyuk**: Writing – review & editing, Supervision, Resources, Methodology, Steven L. Taylor: Writing – review & editing, Writing – original draft, Visualization, Supervision, Resources, Methodology, Investigation, Funding acquisition, Formal analysis, Conceptualization. **Luke Grundy:** Writing – review & editing, Writing – original draft, Visualization, Supervision, Resources, Methodology, Investigation, Funding acquisition, Formal analysis, Conceptualization.

## Supplementary Figure legends

**Figure S1.**
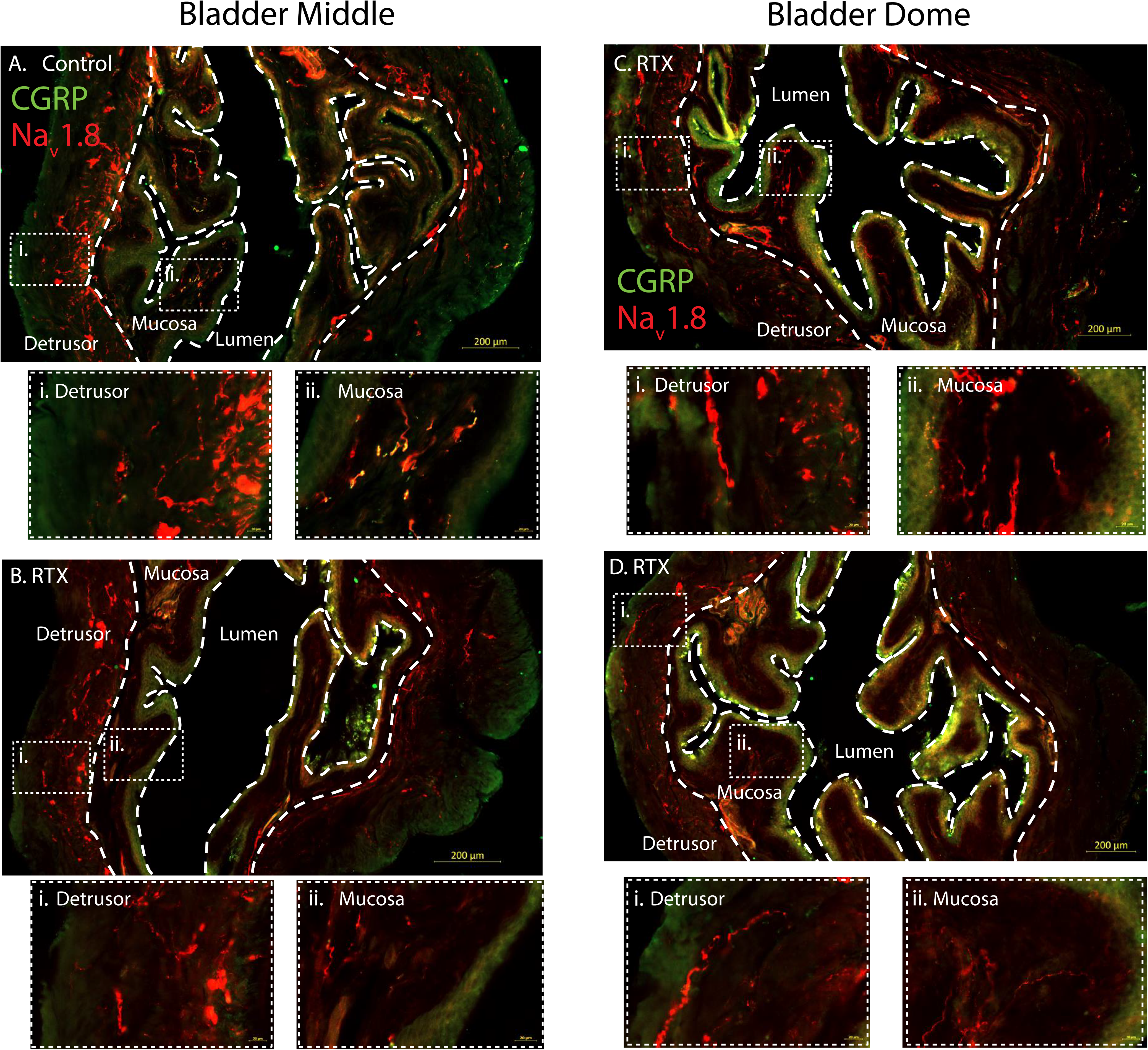
Labelling of Na_v_1.8^-Cre-tdTomato^ bladder with calcitonin gene-related peptide (CGRP) from control- and resiniferatoxin (RTX)-treated mice. Representative photomicrographs showing the distribution of CGRP-immunoreactive fibres (green) and Na_v_1.8-tdTomato expressing fibres (red) in cross-sections of the bladder from the middle (**A** and **B**) and dome (**C** and **D**) regions of control and RTX-treated mice, respectively. Fibres were located within the detrusor and the mucosa. High magnification of the afferent fibres within the detrusor (i) and mucosa (ii) are indicated by dashed boxes. Scale bar = 200µm (overall) and 20µm (inserts).

**Figure S2.**
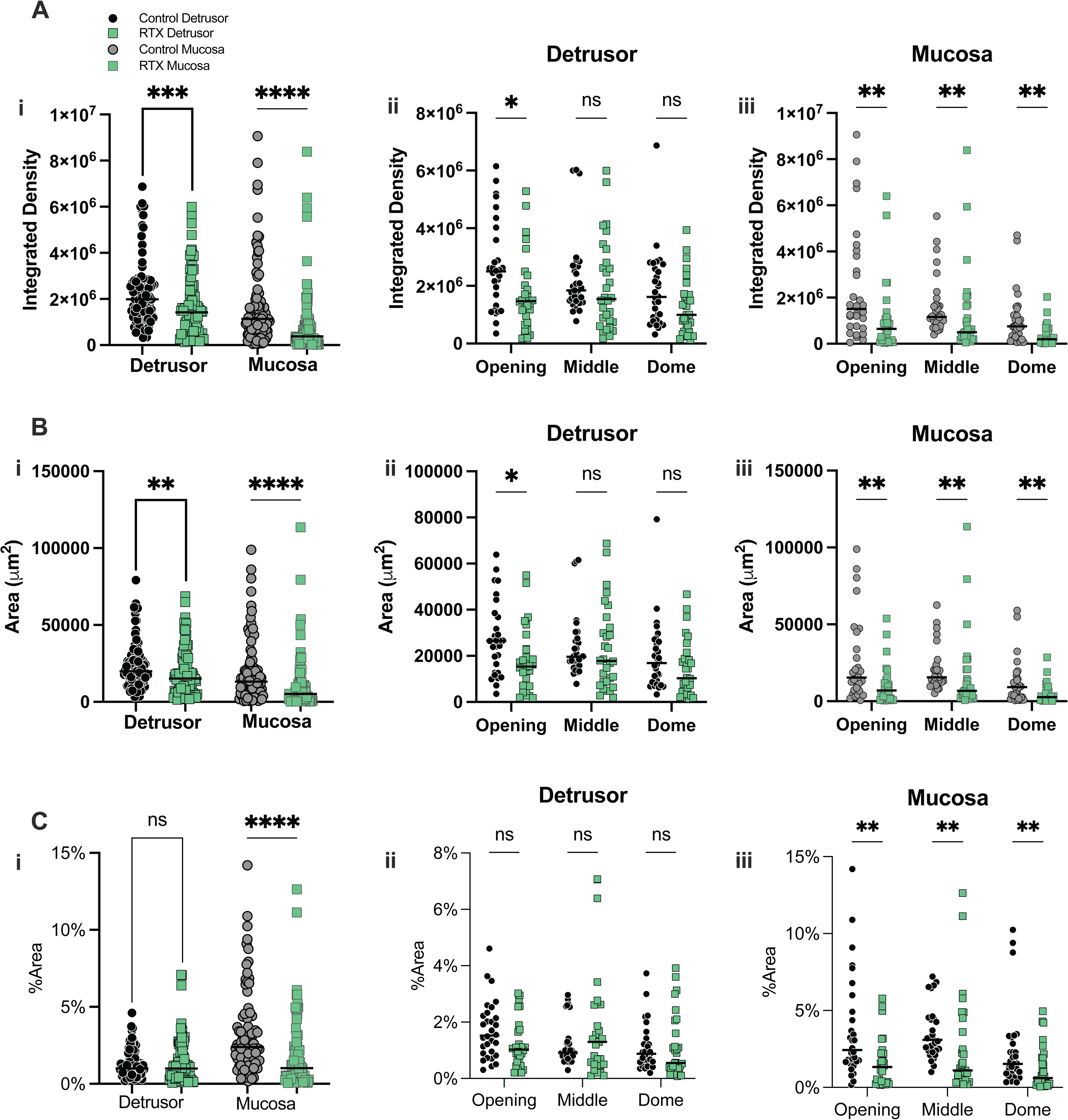
Integrated density, area and %area measurements of calcitonin gene-related peptide (CGRP) fluorescence within control and resiniferatoxin (RTX)-treated bladders. Quantitative data from cross-sections of bladders from control and RTX-treated Na_v_1.8-TdTom mice showing CGRP-AF488 fluorescence (represented in Fig 1, Fig S1) comparing the (A) integrated density, (B) area and (C) %area between control and RTX treated mice (green) in the detrusor (black data points) and mucosa (grey data points) across the (i) entire bladder, and at the bladder opening, middle and dome of the (ii) detrusor and (iii) mucosa layer. Individual data points represent sections (n = 90) from N = 5 mice/group. Data is presented as the median of individual values and analysed using non-parametric Mann-Whitney test, where *P < 0.05, **P < 0.01, ***P < 0.001 and ****P < 0.0001.

**Figure S3:**
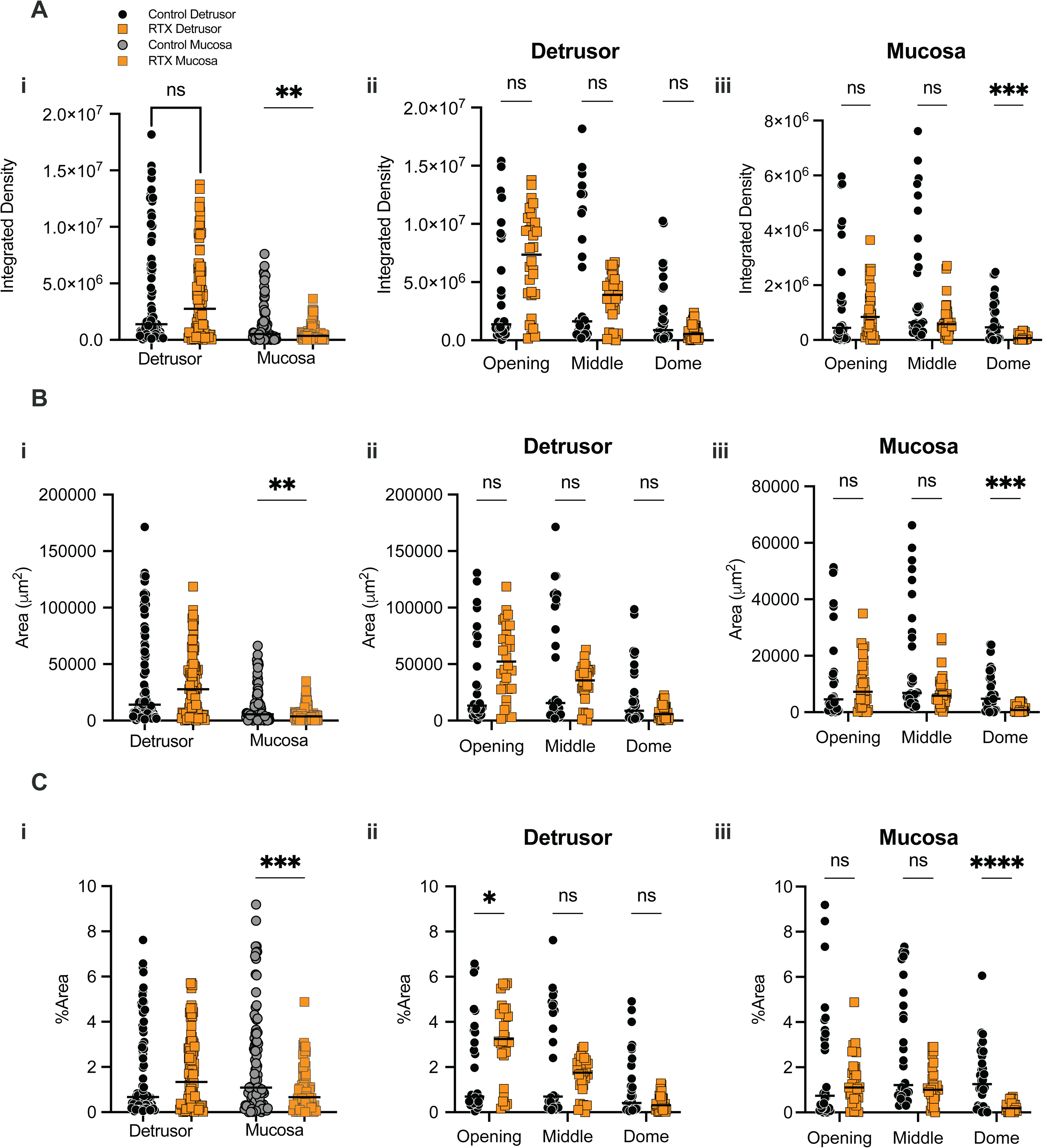
Integrated density, area and %area measurements of Na_v_1.8-tdTomato fluorescence within control and resiniferatoxin (RTX)-treated bladders. Quantitative data from cross-sections of bladders from control and RTX-treated Na_v_1.8-TdTom mice showing TdTom fluorescence (represented in Fig 1, Fig S1) comparing the (A) integrated density, (B) area and (C) %area between control and RTX treated mice (orange) in the detrusor (black data points) and mucosa (grey data points) across the (i) entire bladder, and at the bladder opening, middle and dome of the (ii) detrusor and (iii) mucosa layer. Individual data points represent the same sections (n = 90) from N = 5 mice/group as in Fig 1, Fig S2. Data is presented as the median of individual values and analysed using non-parametric Mann-Whitney test, where *P < 0.05, **P < 0.01, ***P < 0.001 and ****P < 0.0001.

**Figure S4.**
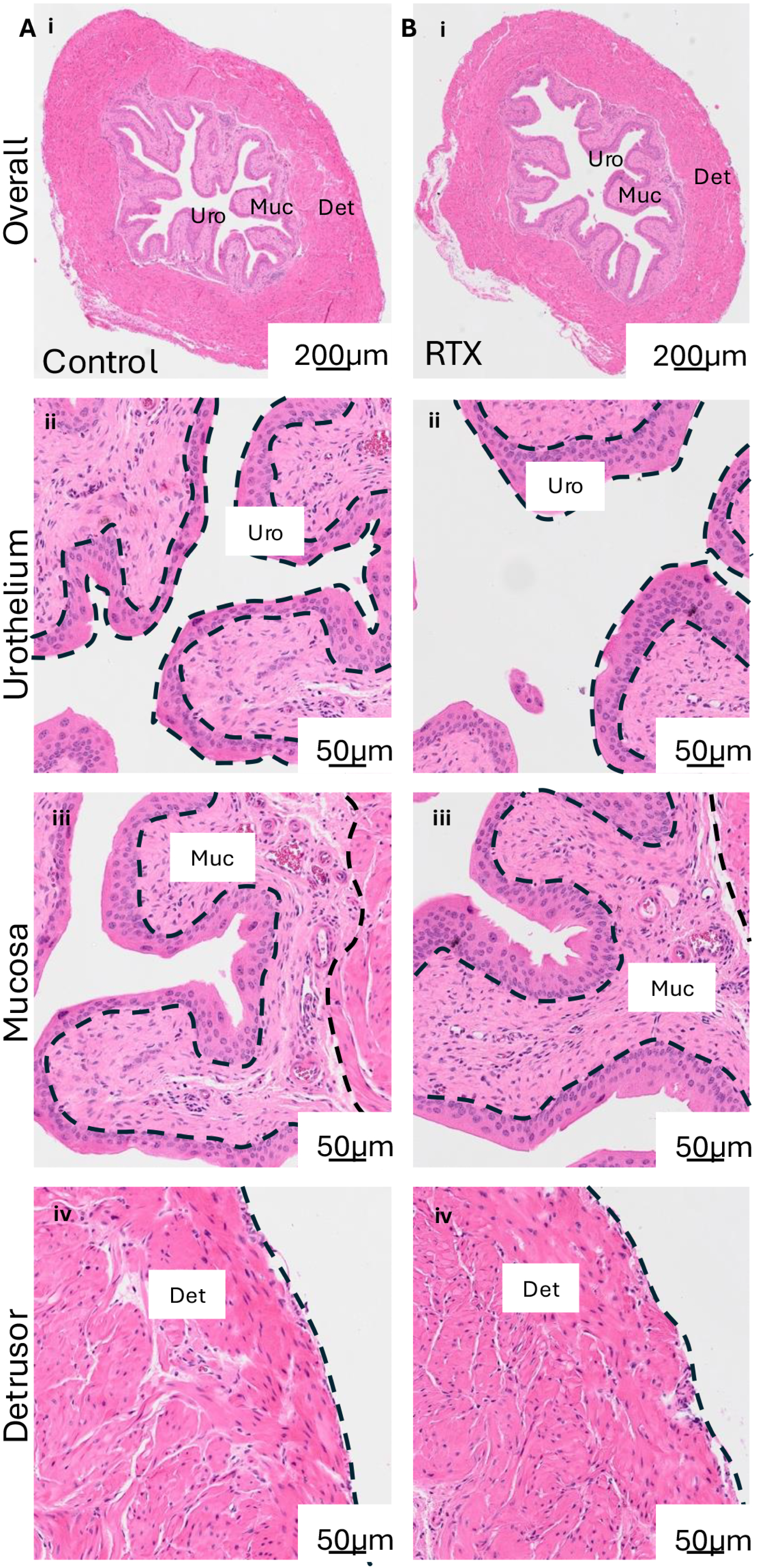
Representative images of hematoxylin and eosin-stained bladder sections from control and RTX-treated mice. Bladders from (**A**) control and (**B**) RTX-treated mice were fixed in 4% PFA and paraffin-sectioned at 5mm. Sections underwent hematoxylin and eosin staining to bladder tissue integrity. Closeups of the (i) urothelium, (ii) mucosa and (iii) detrusor of the control and RTX bladders. No obvious difference in morphology was observed between control and RTX-treated bladders, including intact urothelium.

**Figure S5:**
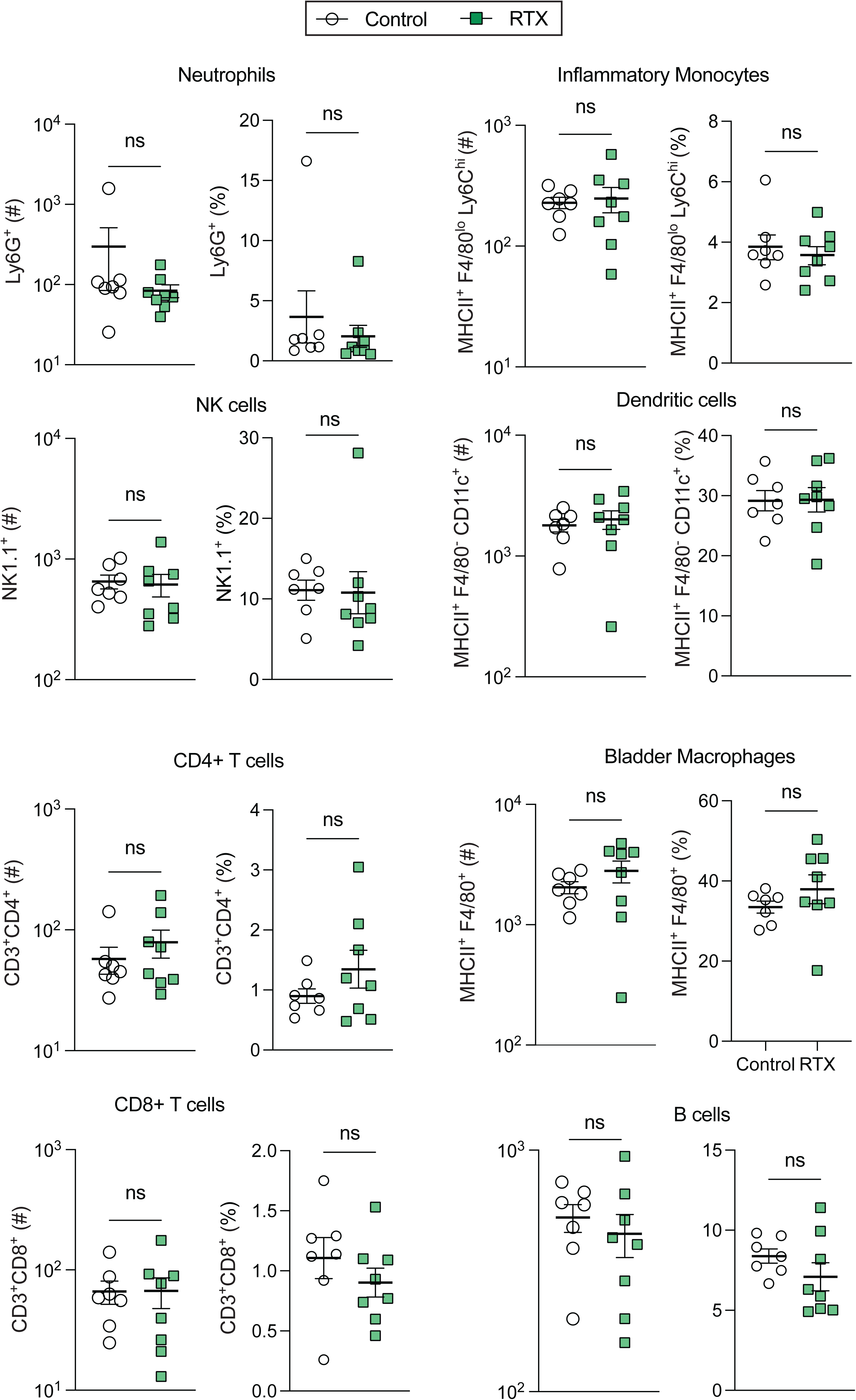
Quantification of immune cells from control and resiniferatoxin (RTX)-treated bladders. Bladder immune cells were sorted and quantified using flow cytometry and the gating strategy outlined in Fig S12. Neutrophils (Ly6G^+^), Inflammatory monocytes (MHCII^+^ F4/80^lo^ Ly6C^hi^), NK cells (NK1.1^+^), Dendritic cells (MHCII^+^ F4/80^-^ CD11c^+^), CD4^+^ T cells (CD3^+^CD4^+^), bladder macrophages (MHCII^+^ F4/80^+^), CD8^+^ T cells (CD3^+^CD8^+^), and B cell (CD19^+^) populations seven days after RTX treatment (green data points) compared to control (black data points) bladders. Individual points represent cell counts from an individual mouse, N = 8 mice. Data was presented as mean ± SEM. Data were compared using Mann-Whitney test for neutrophils and CD4^+^ T cells datasets, and Welch’s t-test for bladder macrophages, inflammatory monocytes, B cells, dendritic cells, NK cells and CD8^+^ T cells datasets.

**Figure S6.**
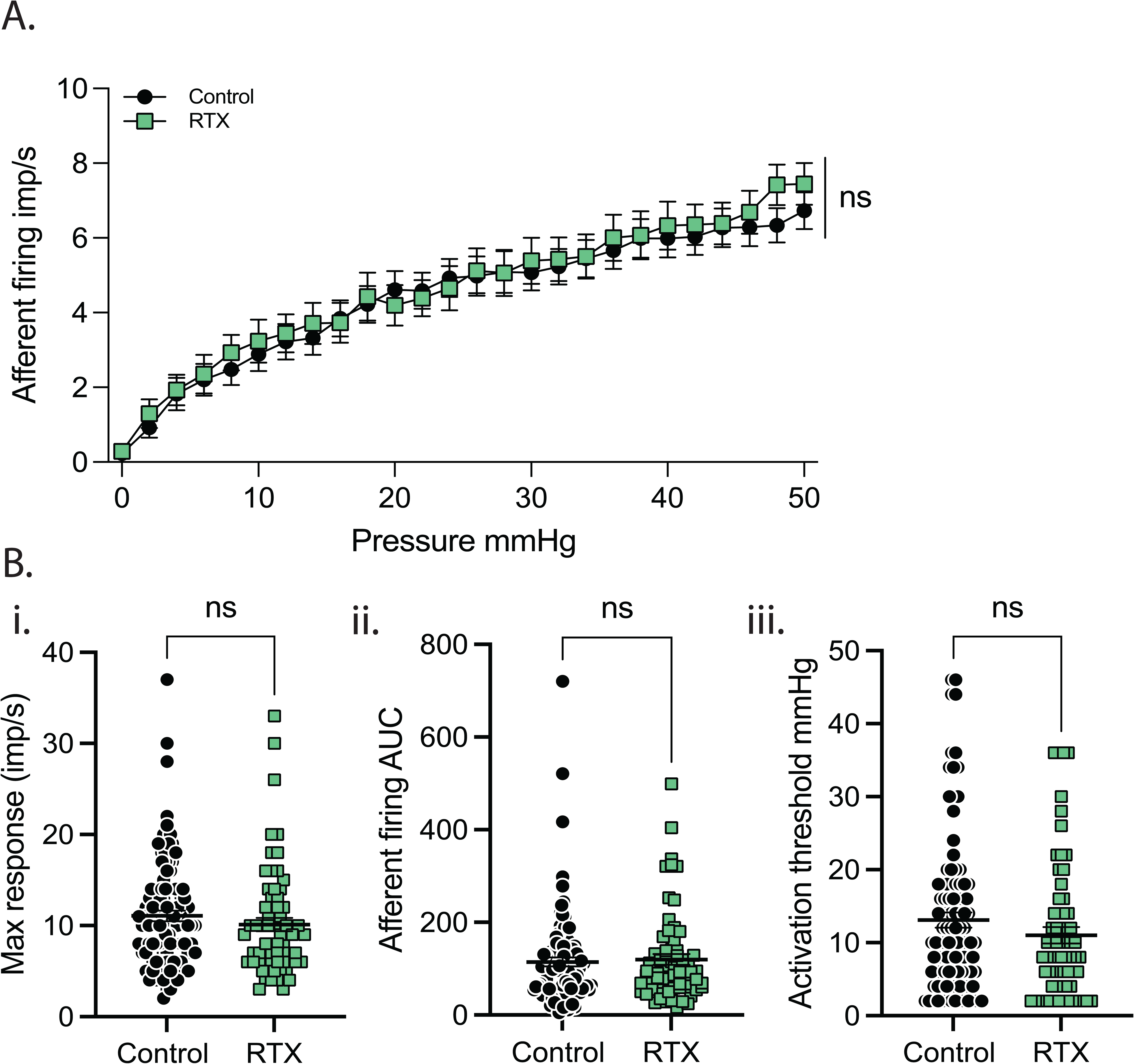
Resiniferatoxin (RTX) treatment does not affect single-unit bladder afferent mechanosensitive responses to distension. (**A**) Comparative data showing single-unit sensitivity to graded distension (0-50 mmHg) between control (black) and RTX (green) treated bladders from Fig 3 (n=7-21 units per experiment from N=5 mice/group). (**B**) (**i**) Max responses, (**ii**) total area under the curve (AUC) of the afferent response to distension and (**iii**) activation thresholds (mmHg) of individual bladder afferent units were not changed by RTX pretreatment compared to control. Data are represented as mean ± SEM, individual data points in (**Bi-iii**) represent values collected from each single unit. Data were compared using two-way ANOVA with Šídák multiple comparisons for A, and unpaired Mann-Whitney test for B.

**Figure S7.**
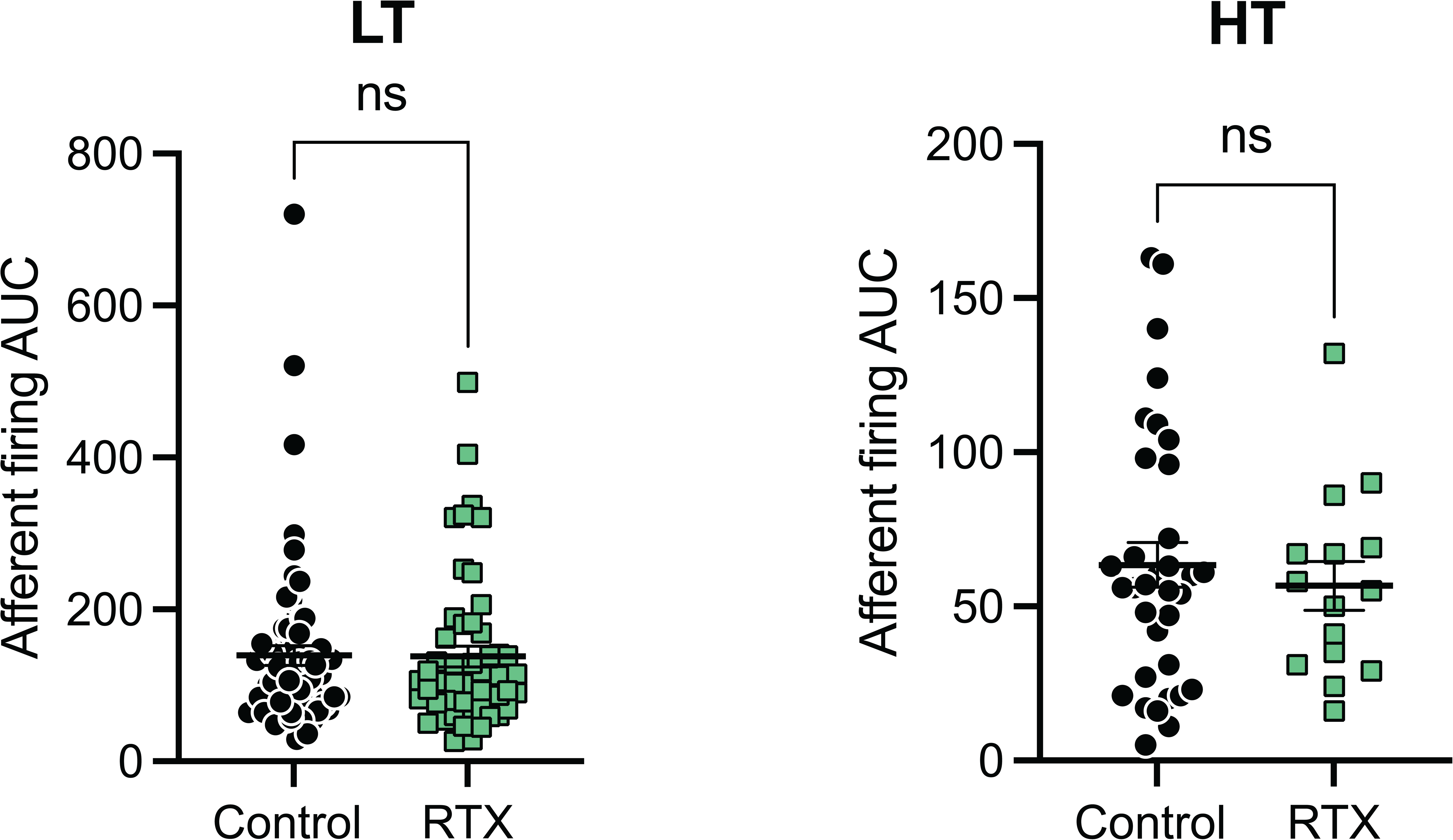
Resiniferatoxin (RTX) does not affect low- or high-threshold mechanosensitive bladder afferent area under the curve. Total area under the curve (AUC) of the afferent response to distension (0-50mmHg) of individual low- (LT) and high- (HT) threshold bladder afferent units in control and RTX treated mice from Fig 3. Data are represented as mean ± SEM, individual data points represent values collected from each single unit. Data were compared using unpaired Mann-Whitney test.

**Figure S8.**
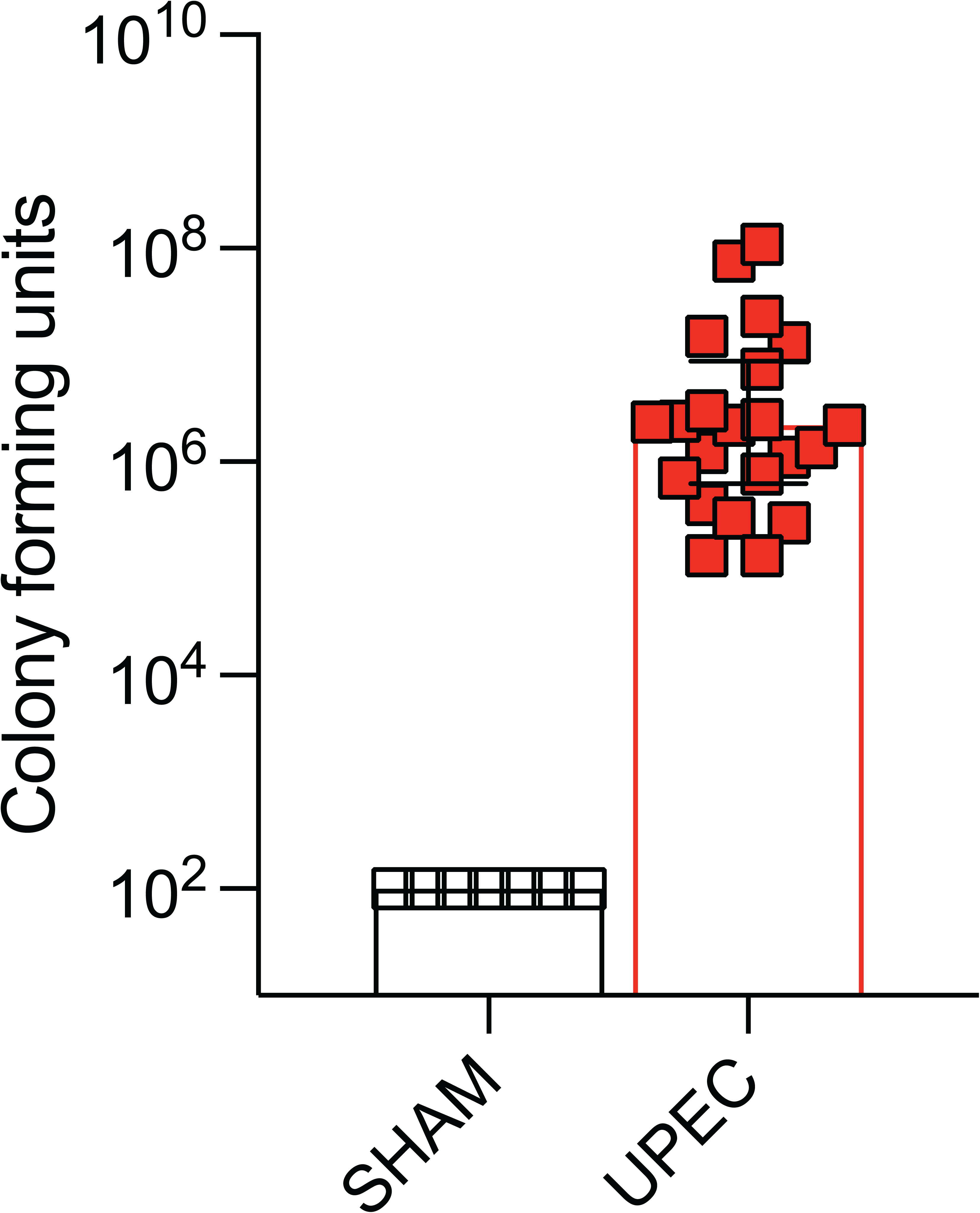
Urine CFU 24hrs after UPEC or Sham treatment. UPEC CFU in urine 24hrs after transurethral bladder inoculation with either PBS (Control N=6) or UPEC strain CFT073 (N=21). Bacteria were found in the urine of UPEC but not sham treated mice.

**Figure S9.**
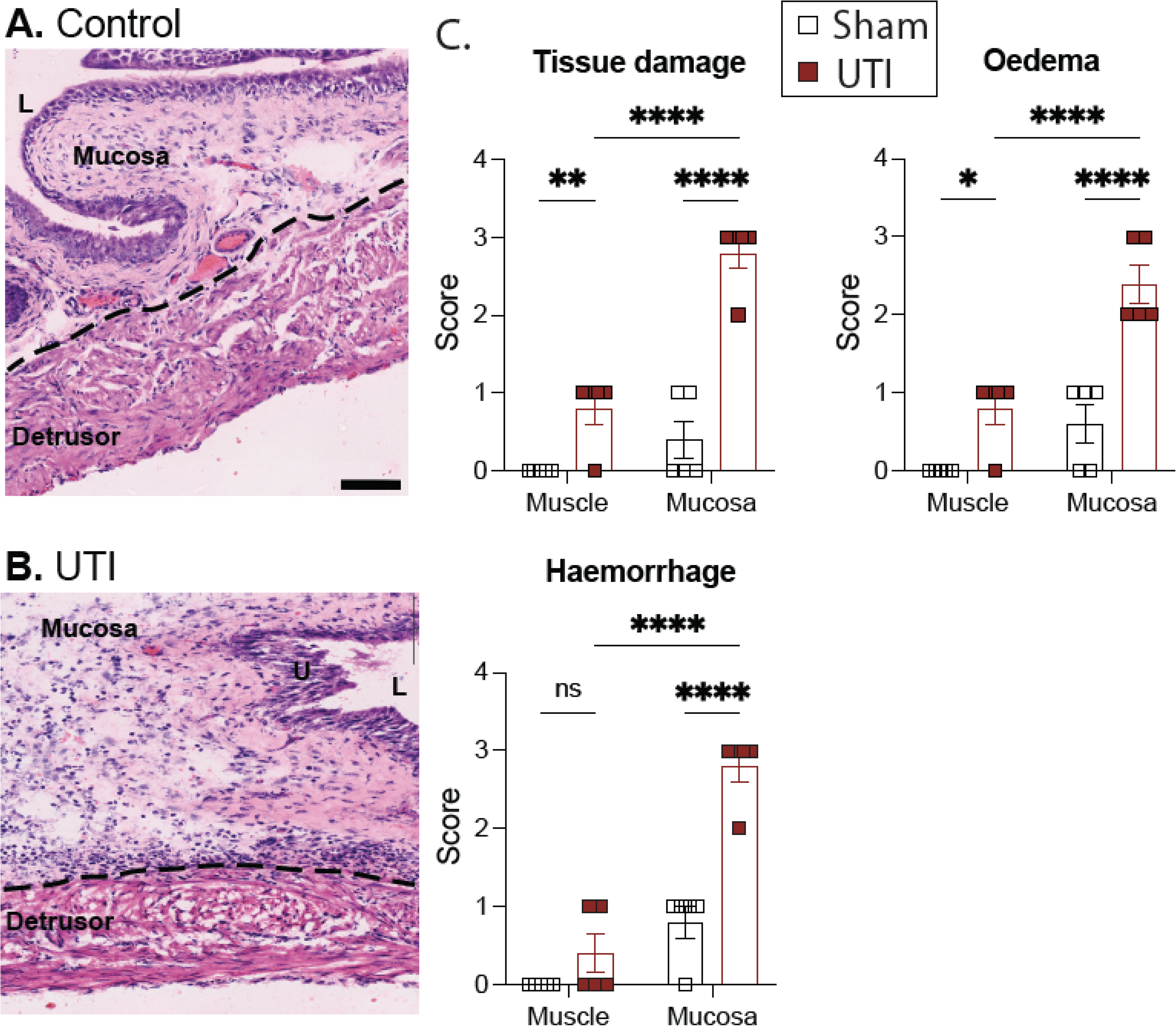
UPEC bladder infection evokes significant bladder damage which predominates in the mucosa. H&E-stained bladder sections from mice 24hrs after sham **(A)** or UPEC instillation **(B)**. **(C)** Histological quantification of bladder damage (using scoring criteria detailed in Table 2) demonstrates a significant increase in tissue damage, oedema, and haemorrhage, in bladders from UTI treated mice relative to Sham-treated mice. Scale bar=100mm. Data are presented as mean ± SEM and analysed by one-way ANOVA (n=5/group). **Figure S10.**

**Supplementary Figure 10.**
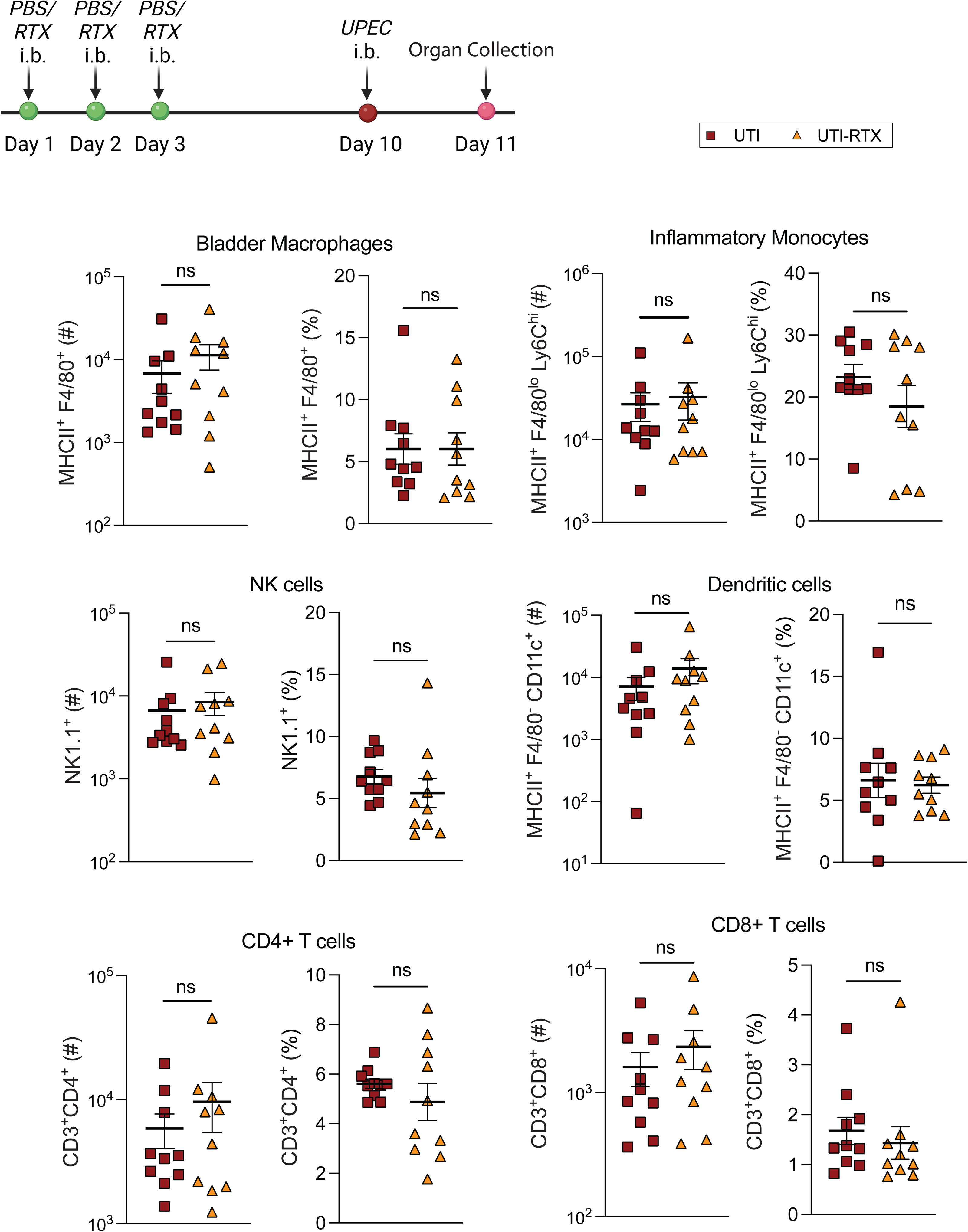
Quantification of immune cells from UTI and resiniferatoxin (RTX)-UTI bladders. Bladder immune cells were sorted and quantified using flow cytometry and the gating strategy outlined in Fig S12. Macrophages (MHCII^+^ F4/80^+^), Inflammatory monocytes (MHCII^+^ F4/80^lo^ Ly6C^hi^), NK cells (NK1.1^+^), Dendritic cells (MHCII^+^ F4/80^-^ CD11c^+^), CD4^+^ T cells (CD3^+^CD4^+^) and CD8^+^ T cells (CD3^+^CD8^+^), from bladders one day after UPEC instillation in mice pretreated with RTX (orange data points) compared to UPEC only (red data points). Individual points represent cell counts (#) and cell frequency (%) from an individual mouse, N = 10 mice. Data was presented as mean ± SEM, Data were analysed by the non-parametric Mann-Whitney test for all datasets.

**Figure S11.**
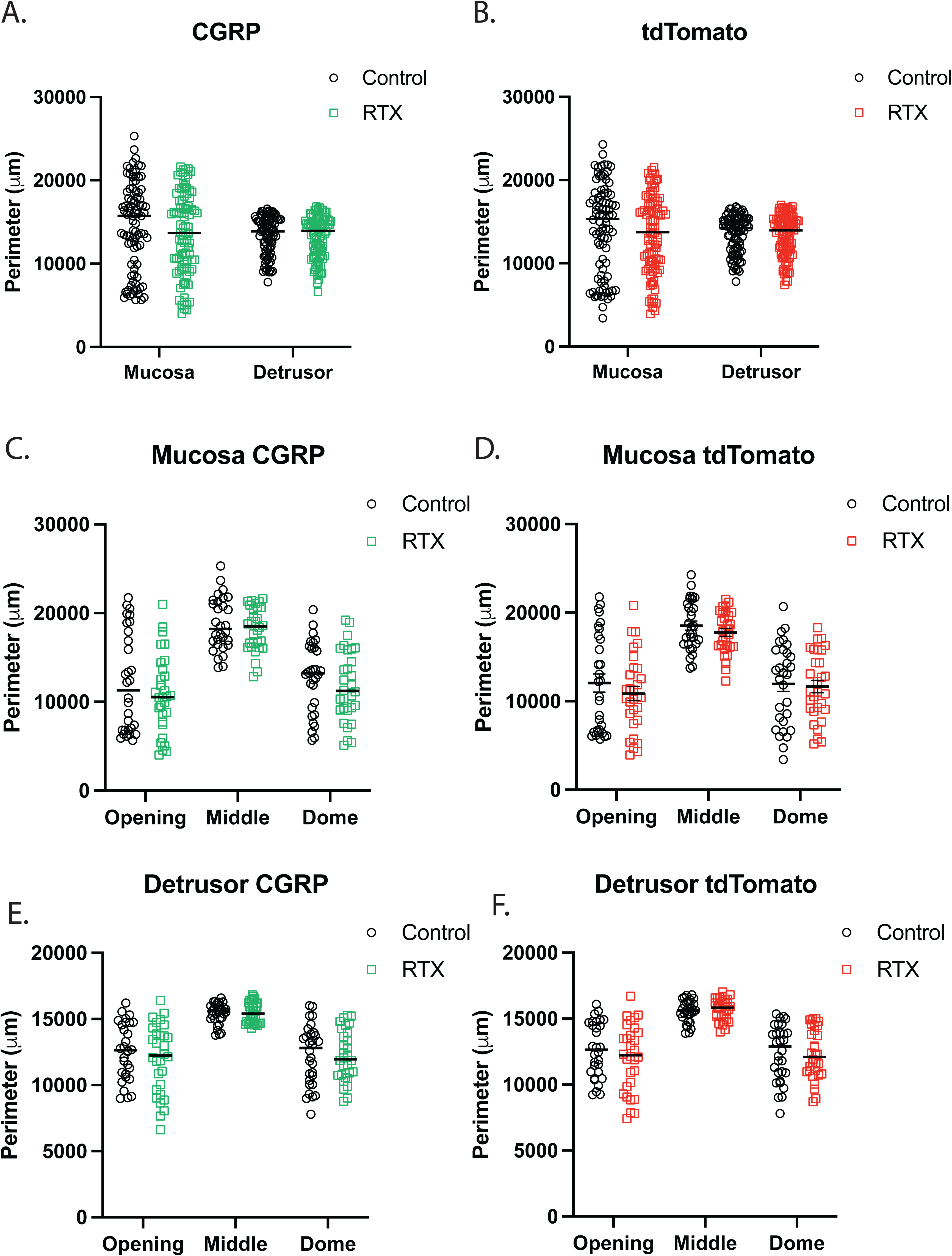
(in methods): Perimeter distance of control and RTX mucosa and detrusor bladder layers. Bladders from control and RTX-treated Na_v_1.8^-Cre-tdTomato^ mice were immunostained with CGRP were sectioned and imaged using the Zeiss Axio Scan.Z1 slide scanner. A total of n=90 sections from N=5 bladders/group were measured for perimeter of mucosa and detrusor using ImageJ. Individual measurements from the detrusor and mucosa were compared overall and at the opening, middle and dome of the bladder. This data was plotted to demonstrate the consistency in selecting areas within the detrusor and mucosa across all samples. The median and ranges was similar across all graphs: (A) CGRP overall, (B) tdTomato overall, (C) mucosa CGRP, (D) mucosa tdTomato, (E) detrusor CGRP and (F) detrusor tdTomato.

**Figure S12.**
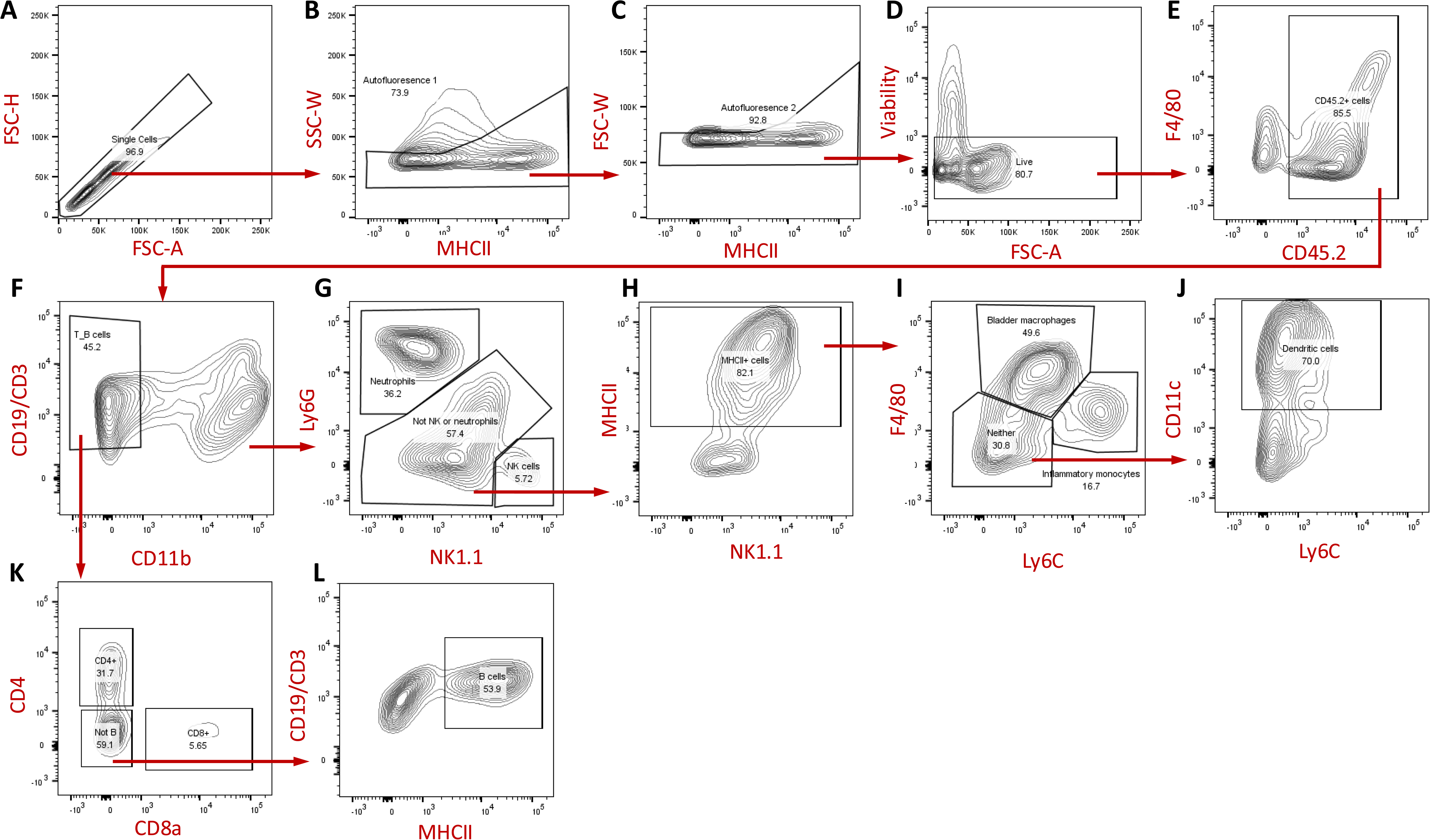
(In methods): Gating strategy for flow cytometry of immune cells in the bladder. (A) Forward scatter height (FSC-H) versus forward scatter area (FSC-A) plot is used to select single cells. (B) Side scatter width (SSC-W) versus MHCII to exclude autofluorescent cells, and further gating on MHCII+ cells is applied. (C) FSC-W versus MHCII for size exclusion of dead cells and debris. (D) Viability stain used to gate live cells. (E) Gating for CD45.2+ cells as a marker of immune cells. (F) CD19 and CD3 are used to gate out B cells (CD19+) and T cells (CD3+). CD11b is used to gate CD11b+ cells. (G) Ly6G and NK1.1 are used to distinguish neutrophils (Ly6G+), NK cells (NK1.1+), and other cell populations (Ly6G-/NK1.1-). (H) MHCII versus NK1.1 gating helps to further refine NK cells and other populations. (I) F4/80 and Ly6C are used to differentiate bladder macrophages (F4/80+), inflammatory monocytes (Ly6C+), and neutrophils. (J) CD11c and Ly6C gating identifies dendritic cells (CD11c+) and monocytes (Ly6C+). (K) CD4 versus CD8a is used to distinguish CD4+ helper T cells and CD8+ cytotoxic T cells among CD3+ populations. (L) MHCII versus CD19/CD3 shows MHCII+ cells with the CD19/CD3- population to identify antigen-presenting cells.

